# Privacy Preserving RNA-Model Validation Across Laboratories

**DOI:** 10.1101/2021.04.01.437893

**Authors:** Talal Ahmed, Mark A Carty, Stephane Wenric, Jonathan R Dry, Ameen Abdulla Salahudeen, Aly A. Khan, Eric Lefkofsky, Martin C Stumpe, Raphael Pelossof

## Abstract

Reproducibility of results obtained using RNA data across labs remains a major hurdle in cancer research. Often, molecular predictors trained on one dataset cannot be applied to another due to differences in RNA library preparation and quantification. While current RNA correction algorithms may overcome these differences, they require access to all patient-level data, which necessitates the sharing of training data for predictors when sharing predictors. Here, we describe SpinAdapt, an unsupervised RNA correction algorithm that enables the transfer of molecular models without requiring access to patient-level data. It computes data corrections only via aggregate statistics of each dataset, thereby maintaining patient data privacy. Furthermore, SpinAdapt can correct new samples, thereby enabling evaluation of validation cohorts. Despite an inherent tradeoff between privacy and performance, SpinAdapt outperforms current correction methods that require patient-level data access. We expect this novel correction paradigm to enhance research reproducibility and patient privacy. Finally, SpinAdapt lays a mathematical framework that can be extended to other -omics modalities.

## Introduction

The advent of high-throughput gene expression profiling has powered the development of sophisticated molecular models to capture complex biological patterns. To ensure the generalization of molecular patterns across independent studies, molecular predictors require validation across platforms and laboratories. However, the transfer of predictors across laboratories still remains a technical obstacle. Batch-specific effects that dominate the biological signal exist between different technologies, laboratories and even library preparation protocols within the same laboratory ^1^. Furthermore, often these inter-institutional datasets are siloed due to human subject privacy concerns. There is an unmet need for a technology that enables the transfer of molecular predictors across labs in a privacy-preserving manner such that sample-level patient data is not transferred.

Correction of batch-specific biases in RNA-Seq datasets has been an active field of research in the past two decades. Numerous methods are proposed to correct batch effects, and these mostly fall into two categories: batch integration and batch correction. Batch integration entails joint embedding of batch-biased expression data in a shared embedding space where batch variations are minimized ^2,3^. Batch correction removes batch biases in the gene expression space, harmonizing batch-biased dataset(s) to a reference dataset. For batch correction, we refer to the reference dataset as the target and the batch-biased dataset as the source. Since the reference remains unchanged in batch correction, the asymmetry enables the transfer (application) of models trained on reference dataset to batch-corrected dataset(s).

Machine learning models in clinical research are often developed using RNA expression data ^4^. Batch integration methods may not be suitable for external validation of classifiers trained on gene expression datasets, since integration methods do not necessarily output expression profiles. These include methods based on gene-wise linear models like Limma^5^, mutually nearest neighbors (MNNs) like MNN Correct ^6^ and Scanorama ^7^, mutually nearest clusters (MNCs) like ScGen ^8^, pseudoreplicates like ScMerge ^9^, and multi-batch clusters like Harmony ^10^. In contrast, batch correction methods can correct a source library to a target reference library, like Combat ^11^, Seurat3 ^12^ and the proposed SpinAdapt algorithm, and thus can be used for transferring classifiers across expression datasets.

Prior integration and correction methods require full sample-level access for integration or correction of datasets. Therefore, the transfer of molecular predictors between laboratories necessitates the transfer of patient-level training data for the molecular predictor. This data access requirement can inhibit the transfer of models between laboratories, given transfer of data may not be possible due to data ownership, GDPR, or similar regulations. To overcome these challenges, we present SpinAdapt, a method that enables the transfer of molecular predictors between laboratories without disclosure of the sample-level training data for the predictor, thereby allowing laboratories to maintain ownership of the training data and protect patient privacy. Instead of sharing sample-level data, privacy-preserving aggregate statistics of the training data are shared along with the molecular predictor. Our approach is based on the concept of matrix masks from privacy literature, where the sample-level data and matrix mask is kept private, while the output of the matrix mask is shared publicly.

This study demonstrates the transfer and validation of diagnostic and prognostic models across transcriptomic datasets, using SpinAdapt, while drawing comparisons with other batch correction methods. The common task of integration (homogenization) across multiple transcriptomic datasets is also evaluated for multiple cancer types, comparing various integration methods. In our experiments, SpinAdapt outperforms other batch correction methods in the majority of these diagnostic, prognostic, and integration tasks, without requiring direct access to sample-level data. Therefore, SpinAdapt may also be preferable for sharing molecular predictors across labs where the training dataset can be shared and data privacy is not an issue.

## Results

### Algorithm overview

We aimed to develop a framework for transfer and validation of molecular predictors across platforms, laboratories, and varying technical conditions. Furthermore, we aimed to remove the requirement of sharing training data in order to evaluate and validate predictors across labs. To this end, we developed SpinAdapt which enables validation of predictors while preserving data privacy (**Figure 1A)**. Data factors, which are aggregate statistics of each dataset, neither convey Protected Health Information (PHI) nor allow reconstruction of sample-level data (**supplementary note**), and thus can be shared externally. SpinAdapt learns corrections between data factors of each dataset, followed by application of corrections on the biased expression dataset (source). Note that our framework enables the correction of new data samples, which has important implications as discussed later.

**Figure 1.**
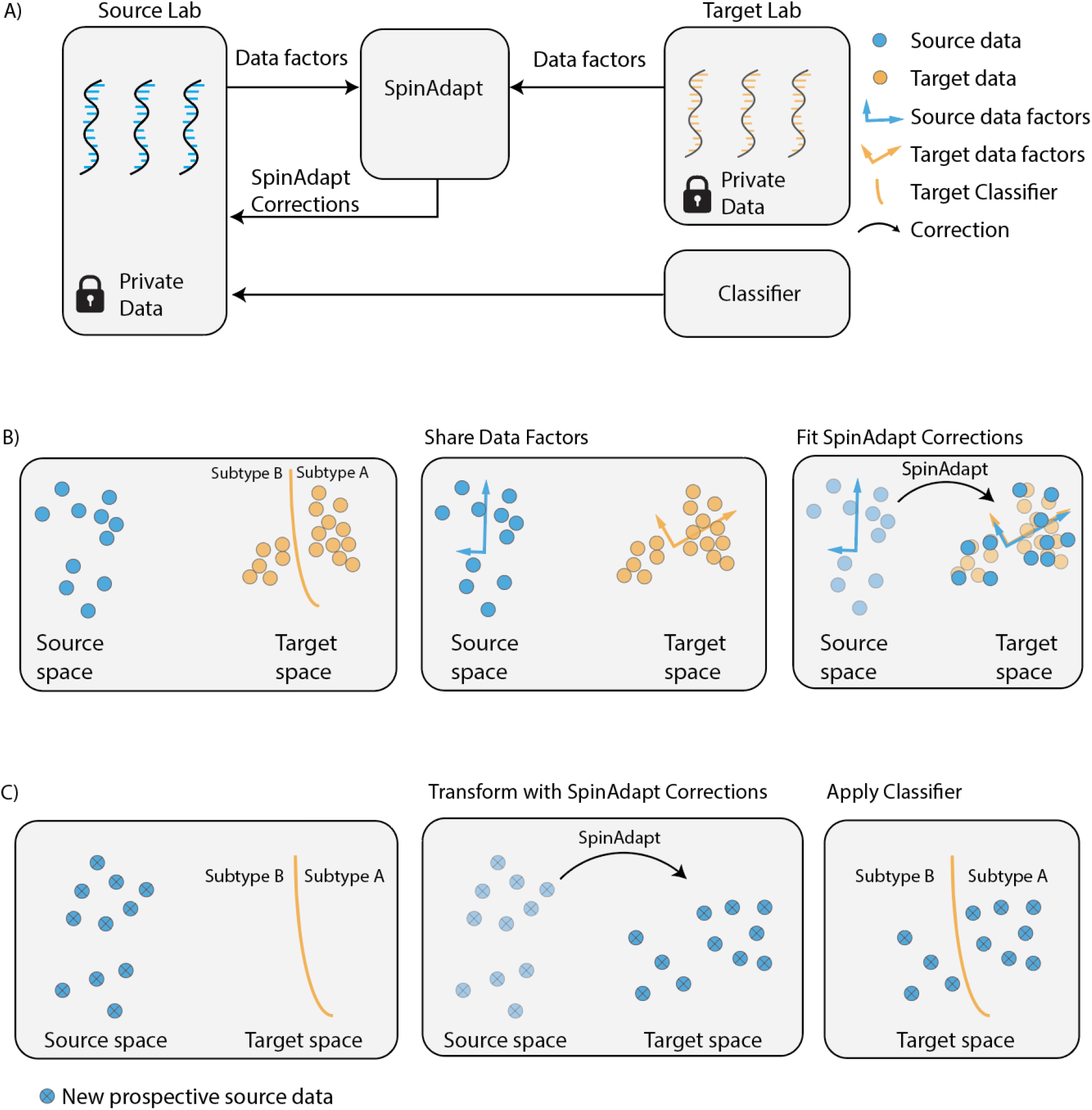
Privacy-preserving transfer of molecular models between a target lab and a source lab. **A**) A target dataset with a trained classifier and protected RNA data provides its privacy-preserving RNA factors and a molecular classifier to SpinAdapt. A source dataset used for validation provides its own privacy-preserving RNA factors to SpinAdapt. Given the factors, SpinAdapt returns a correction model to source, where the source data is corrected. Target classifier without modification can then be validated on source-corrected. **B**) Source and target factors are calculated as the principal components of RNA data. Next, SpinAdapt learns a correction model from source to target eigenvectors (factors). **C**) Evaluation of the SpinAdapt correction model on the held-out prospective source data. Finally, the target-trained classifier is applied to the corrected source data.

SpinAdapt corrections are learnt using a regularized linear transformation between the data factors of source and target, which comprise of the PCA (Principal Component Analysis) basis, gene-wise means, and gene-wise standard deviations of source and target, respectively (**methods, Figure 1B**). The linear transformation is the solution of a non-convex objective function, which is optimized using an efficient computational approach based on projected gradient descent. Once the transformation has been learned, it can be applied on the source dataset for correction, followed by application of the target-trained classifier on the corrected source dataset (**Figure 1C)**. Therefore, the learning module requires access only to the data factors of each dataset to learn the transformation for the source dataset.

Since the learning step (**Figure 1B**) is separated from the transform step (**Figure 1C**), the transform step can be applied to new prospective data that was held-out in the learning step. The ability to transform data held-out from model training is deemed necessary for machine learning algorithms to avoid overfitting, by ensuring the test data for the predictor is not used for training. Including predictor test data in training can lead to information leakage and overly optimistic performance metrics. To avoid this, the evaluation data in transform step (transform) is kept independent of the train data in the learning step (fit). This fit-transform paradigm is extended by SpinAdapt to transcriptomic datasets.

The training step of the algorithm is based on the idea of aligning the PCA basis of each dataset. To demonstrate the concept, we apply SpinAdapt on a transcriptomic dataset of paired patients, employing TCGA-BRCA cohort consisting of 481 breast cancer patients, where RNA was profiled both with RNA-seq and microarray. We assigned the RNA-seq library as target and the microarray as source. The application of SpinAdapt aligns the PCA basis of source to target, which resulted in alignment of embeddings as well as gene expression profiles across the paired datasets **(Figure 2A)**. The paired patients were composed of four cancer subtypes: Luminal A (LumA), Luminal B (LumB), Her2, and Basal. When the corrected source dataset is visualized with the target library in a two-dimensional space with UMAP, we observe each of the four subtypes to be harmonized across the two libraries, as compared with before correction **(Figure 2B)**. The subtype-wise homogenization is achieved without the use of subtype labels in the training step, demonstrating that alignment of basis in the PCA space achieves efficient removal of technical biases in the gene-expression space. We also conduct a simulation experiment to explore SpinAdapt for removal of batch effects, where batch effect is simulated between synthetic datasets (source and target), and the source dataset is corrected using SpinAdapt (see results in **Supplementary note, Supplementary Figure 6**).

**Figure 2.**
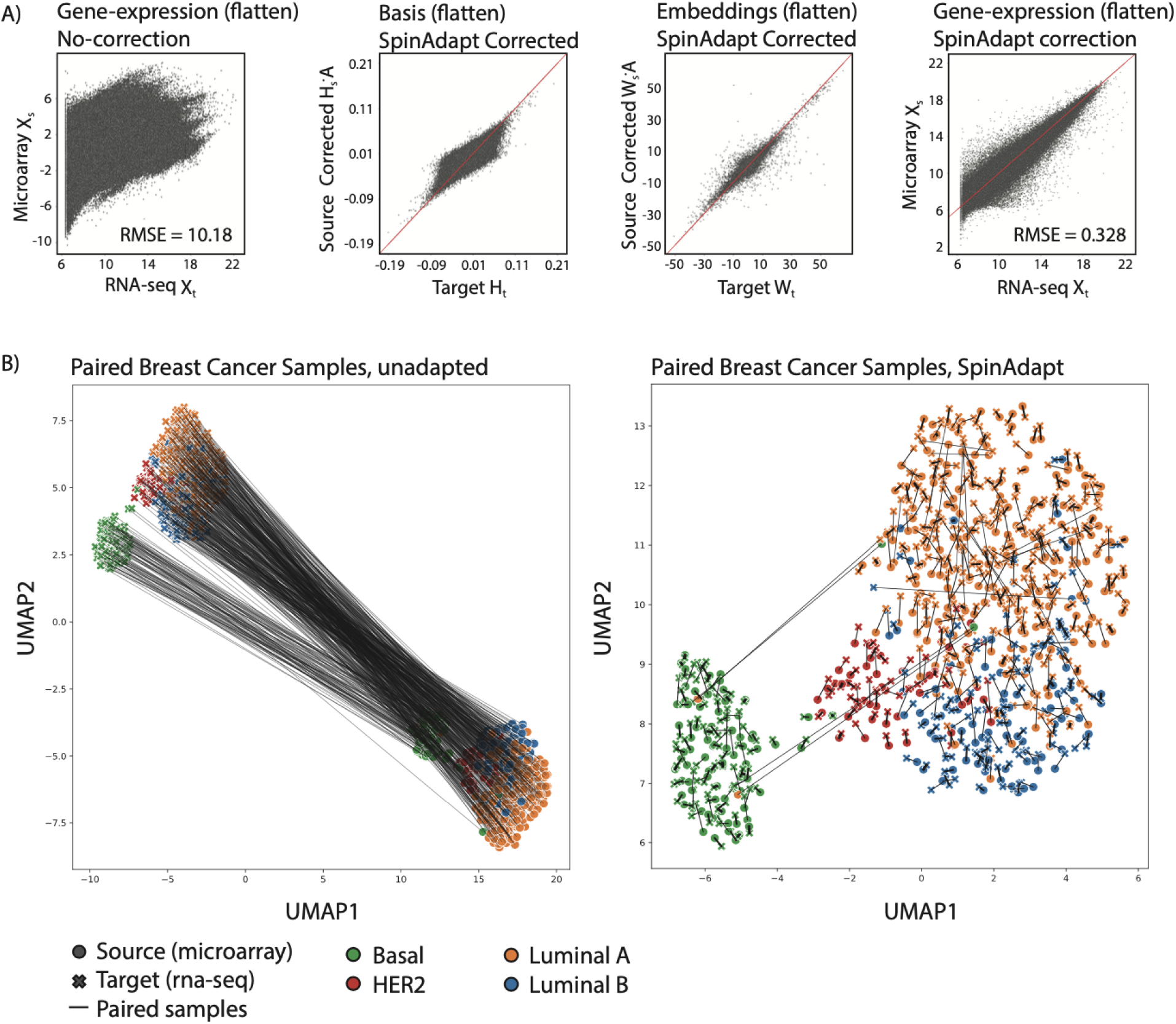
Batch correction performance on paired 481 TCGA-BRCA patients profiled with RNA-seq (target) and microarray (source). **A**) Scatter plots of gene expression values in target with uncorrected source expression, target basis with corrected source basis, target embeddings with corrected source embeddings, and finally gene expression values in target with corrected source expression, where all corrections are performed using SpinAdapt. **B**) Source expression dataset before and after correction, plotted with the reference target library (visualized in 2D with UMAP embeddings). The samples are labeled by cancer subtype and the paired samples are connected with a solid line. Left panel shows cancer subtypes before correction. Right panel shows subtype homogeneity and matching of the paired samples across datasets after SpinAdapt correction.

### Transfer of diagnostic predictors

We demonstrate the transfer of multiple distinct tumor subtype classifiers on four pairs of publicly available cancer datasets (bladder, breast, colorectal, pancreatic), covering 4,076 samples and three technological platforms (RNA-Seq, Affymetrix U133plus2 Microarray, and Human Exon 1.0 ST Microarray) (**Supplementary Table 1**). Cumulatively, we validated the transfer of seventeen tumor subtypes across the four experiments, drawing comparisons of SpinAdapt with other batch correction methods like ComBat and Seurat. For each dataset pair and tumor subtype, we trained a one-vs-rest tumor subtype classifier on the target dataset. The hyperparameters for each subtype classifier were chosen in a cross-validation experiment on the target dataset, while the source dataset was held-out from classifier training (**methods**).

A common approach for validating target-trained classifiers across datasets is to correct all source data to target, and then evaluate the classifier on corrected source dataset. However, such an approach requires the batch correction (adaptation) model to train on the source dataset, which is also the test set for the classifier. Training the adaptation model on the test set may lead to information leakage, which may lead to overly optimistic performance results. The risk of overfitting on the test set has been sparingly discussed in the batch correction literature. We propose a validation framework that holds out a subset of the source data from training of both the adaptation model and subtype classifier. Since the source subset is completely held out from training of both models, it can be used as the test set for the subtype classifier without the risk of information leakage.

Specifically, the validation framework proceeds by creating two mutually exclusive sets from source (Source-A and Source-B). We first fit the adaptation model between Source-A and target, transform Source-B using the adaptation model, followed by prediction on transformed Source-B using the target-trained classifier (**Supplementary Figure 1A**). Similarly, we fit the adaptation model between Source-B and target, followed by transformation and prediction on Source-A (**Supplementary Figure 1B**). Finally, we concatenate the held-out predictions on Source-A and Source-B, followed by performance evaluation using F-1 score (**methods**). SpinAdapt’s performance was evaluated using this framework, so the test set is always held-out from training modules. Existing correction methods, like ComBat and Seurat3, have currently not implemented a transformation method for out-of-sample data that is held-out from their training. Therefore, these methods had to be trained on the classifier test set in the aforementioned framework (**methods**).

We repeated the above experimental framework 30 times and reported the mean F-1 score for each tumor subtype. SpinAdapt significantly outperformed Seurat3 on seven out of the seventeen tumor subtypes including Pancreatic subtypes: Progenitor, ADEX, Immunogenic, Colorectal subtypes: CMS3, Breast subtypes: Luminal A, Bladder subtypes: Squamous and Stroma. SpinAdapt also significantly outperformed ComBat on eleven out of the seventeen subtypes including Pancreatic subtypes: Progenitor, ADEX, Immunogenic, Colorectal subtypes: CMS1, CMS2, CMS3, Breast Subtypes: Her2, Bladder subtypes: Lump P, Lum U, Lum NS, and Stroma. SpinAdapt was not significantly outperformed by either Seurat or ComBat for any subtype (**Figure 2A-D, Supplementary Figure 2, Supplementary Tables 2 and 4, methods**).

### Integration analysis

Dataset integration, an RNA-homogenization task that requires access to sample-leve data, is commonly adopted for single-cell RNA homogenization. To evaluate the tradeoff between privacy preservation and full data access, we compared SpinAapat to Seurat, Combat, Limma^5^, and Scanorama^7^ for integration of bulk-RNA datasets in the four dataset pairs used previously (**Supplementary Table 1**). For high integration performance, we want to maximize dataset mixing while maintaining subtype-wise separability (no mixing of tumor subtypes) within integrated datasets.

To evaluate the various integration methods, we employ Uniform Manifold Approximation and Projection (UMAP) transform in conjunction with the average silhouette width (ASW) and local inverse Simpson’s index (LISI) (**methods**). For each of the four cancer dataset pairs, the silhouette score is computed for each integrated sample in source and target, and then the average silhouette score is reported across all samples (**methods, Figure 3A, Supplementary Table 3**). Even though SpinAdapt did not have access to sample-level data when learning the transformation between source and target, it significantly outperformed each of the other methods for colorectal cancer and breast cancer (P < 10e-5 and *P*<10e-34, respectively). For pancreatic cancer, SpinAdapt outperformed ComBat, Limma, and Scanorama (*P*<0.05 for each method). For bladder cancer, Scanorama outperformed SpinAdapt (*P* < 10e-5), whereas SpinAdapt outperformed ComBat and Limma (*P* < 10e-6) (**Supplementary Table 5**). Even though the integration performance of SpinAdapt can be improved via direct access to samples, it significantly outperforms most of the competing methods in each experiment.

**Figure 3.**
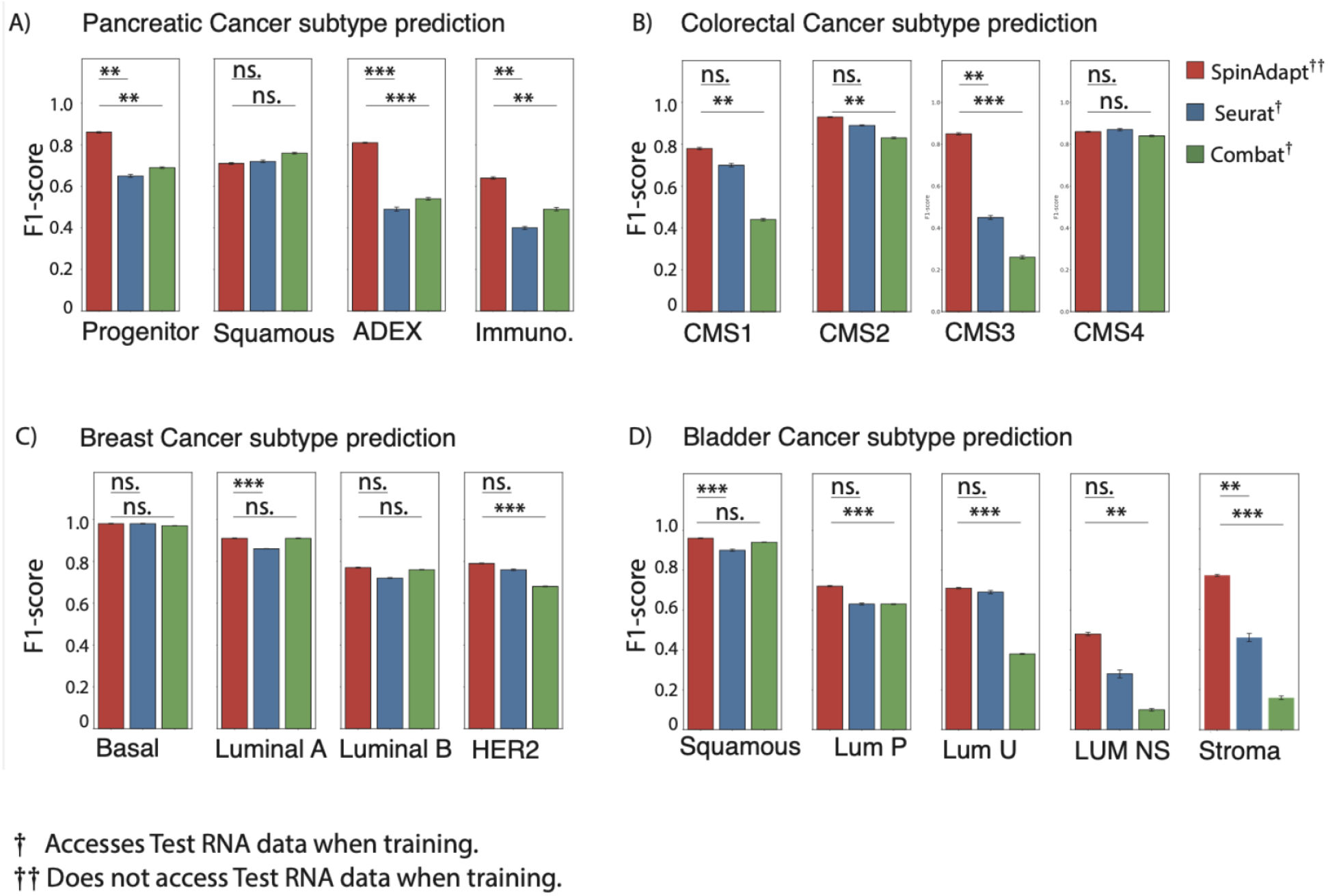
**A-D**) Subtype prediction performance on held-out source subsets. We train subtype predictors on target data and evaluate them on source data. Source data is split into two disjoint subsets such that the correction model is trained on one subset and the predictor performance is evaluated on the other held-out subset. Seurat and ComBat do not support a fit-transform paradigm, and therefore they are trained and evaluated on each of the disjoint subsets. For each subtype, the vertical bar represents the mean F-1 score and the error bar represents the standard error over 30 repetitions of the experiment. SpinAdapt either ties or outperforms Seurat and ComBat on: pancreatic cancer, colorectal cancer, breast cancer, and bladder cancer subtypes. Significance testing by two-sided paired McNemar test (**methods**).

To analyze batch mixing and subtype-wise separation independently, we also employ the LISI metric for integration evaluation. For good integration performance, we sought high batch diversity and low subtype diversity in local sample neighborhoods, which associates with high batch LISI (bLISI) and low tissue LISI (tLISI) score, respectively. For each of the four cancer datasets, we report the average bLISI and tLISI scores across all integrated samples in source and target (**methods, Supplementary Figure 4**).

### Transfer and validation of prognostic predictors

Finally, we demonstrate the transfer of Cox regression models across distinct datasets for three cancer types (breast, colorectal, pancreatic, see **Supplementary Table 5**). Specifically, a Cox Proportional Hazards (PH) model is trained on the target dataset using a gene signature determined through application of an ensemble method on the target dataset (**methods**). The risk thresholds for the survival model are determined based on the upper and lower quartiles of the distributions of log partial hazards of the target dataset, such that samples with a predicted log partial hazard higher than the 75% percentile of said distribution are predicted to be high risk, and samples with a predicted log partial hazard lower than the 25% percentile of said distribution are predicted to be low risk. These model prediction thresholds were fixed across all evaluations.

We then compare multiple batch correction (adaptation) methods for transfer of the prognostic models from target to the source dataset. For each cancer type, the source dataset is adapted to the target using SpinAdapt, Seurat, and ComBat. The target-trained Cox PH model is used to generate predictions (log partial hazards) on all samples from the source dataset, both for the uncorrected source dataset and the three correction methods. The risk thresholds determined on the target dataset are used to classify samples from the source dataset as either low risk, high risk, or unclassified, based on their predicted log partial hazards values. The performance of the prognostic models is quantified by computing the c-indices as well as the 5-year log-rank p-value and 5-year hazard ratio (HR) of the combined predicted high risk and low risk groups of source samples for each cancer type and each adaptation method (**Figure 5, Supplementary Table 7**). SpinAdapt demonstrates high survival prediction accuracy for all datasets (C-index, log-rank p-value, HR): colorectal [0.63, 4e-2, 4.24], breast [0.66, 1e-5, 3.2], pancreatic [0.65, 2e-6, 3.51], in contrast with uncorrected source dataset: colorectal [0.52, 4e-1, 0.54], breast [0.62, 7e-4, 2.2], pancreatic [0.50, 7e-1, 1.29]. Furthermore, SpinAdapt outperforms Seurat: colorectal [0.51, 7e-1, 0.85], breast [0.59, 2e-2, 2.0], pancreatic [0.62, 2e-1, 2.66], as well as ComBat: colorectal [0.56, 2e-1, 0.57], breast [0.61, 3e-4, 2.4], pancreatic [0.62, 7e-5, 2.64], in terms of all performance metrics for all cancer types.

**Figure 4.**
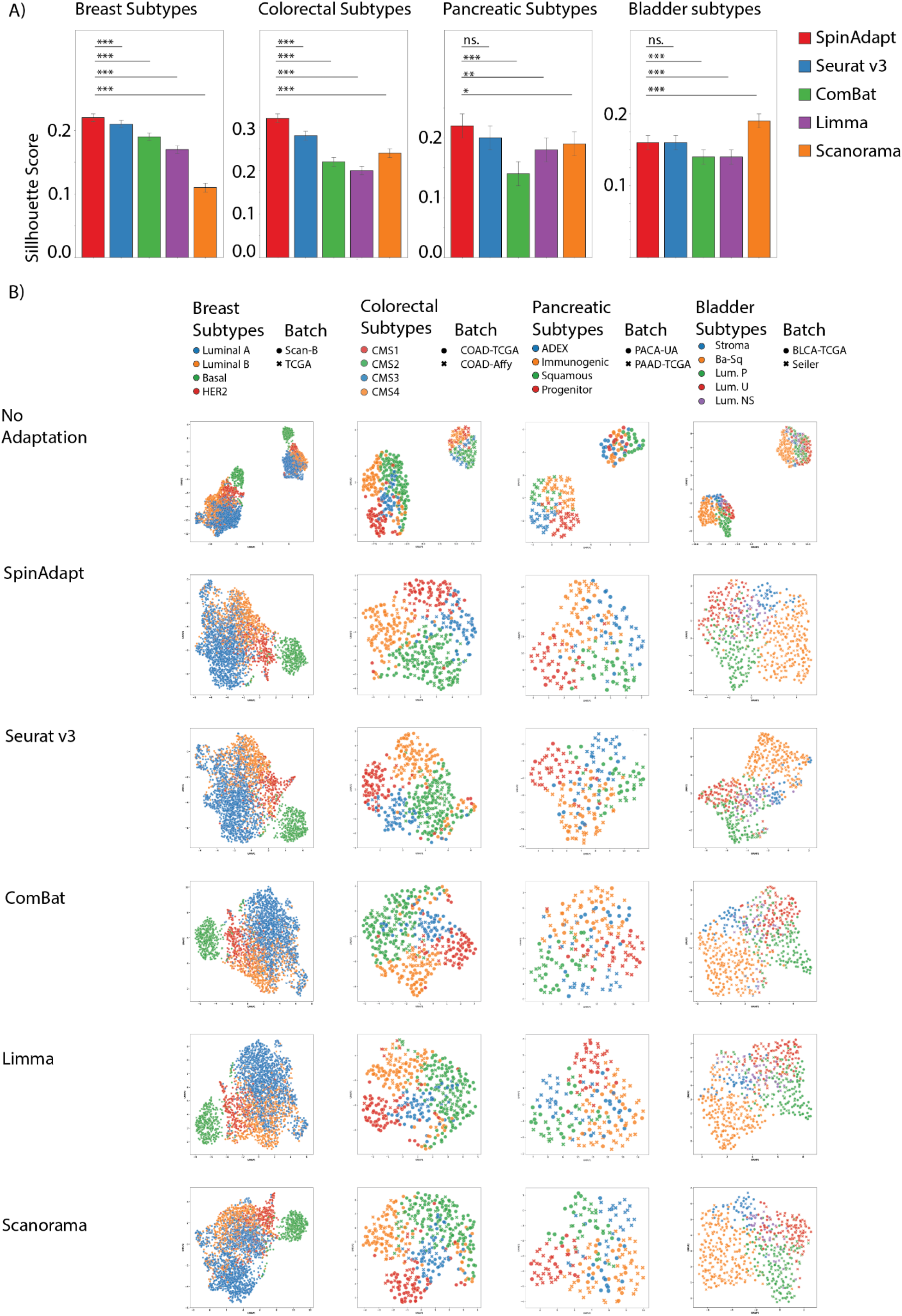
Integration of source with target across the four dataset pairs. **A)** Quantification of integration performance using silhouette score finds SpinAdapt to provide significantly better integrations in breast, colorectal, and pancreatic cancer datasets. Significance testing by two-sided paired Wilcoxon test (**methods**). (ns: *P* ≥ 0.05, \**P* < 0.05, \*\**P* < 0.01, \*\*\**P* < 0.001). Statistical significance is defined at *P* < 0.05. **B)** UMAP plots for dataset integration, labeling samples by cancer subtype. Top panel shows cancer subtypes in each dataset before correction. Subtype homogeneity is apparent in the majority of integration tasks regardless of library size. Subtype mixing is visible in regions where multiple subtypes cluster together. Subtype mixing is observed before and after correction in Breast between luminal subtypes, in Colorectal between CMS 2 and 4, in Pancreatic between ADEX and immunogenic, and in Bladder between luminal subtypes.

**Figure 5.**
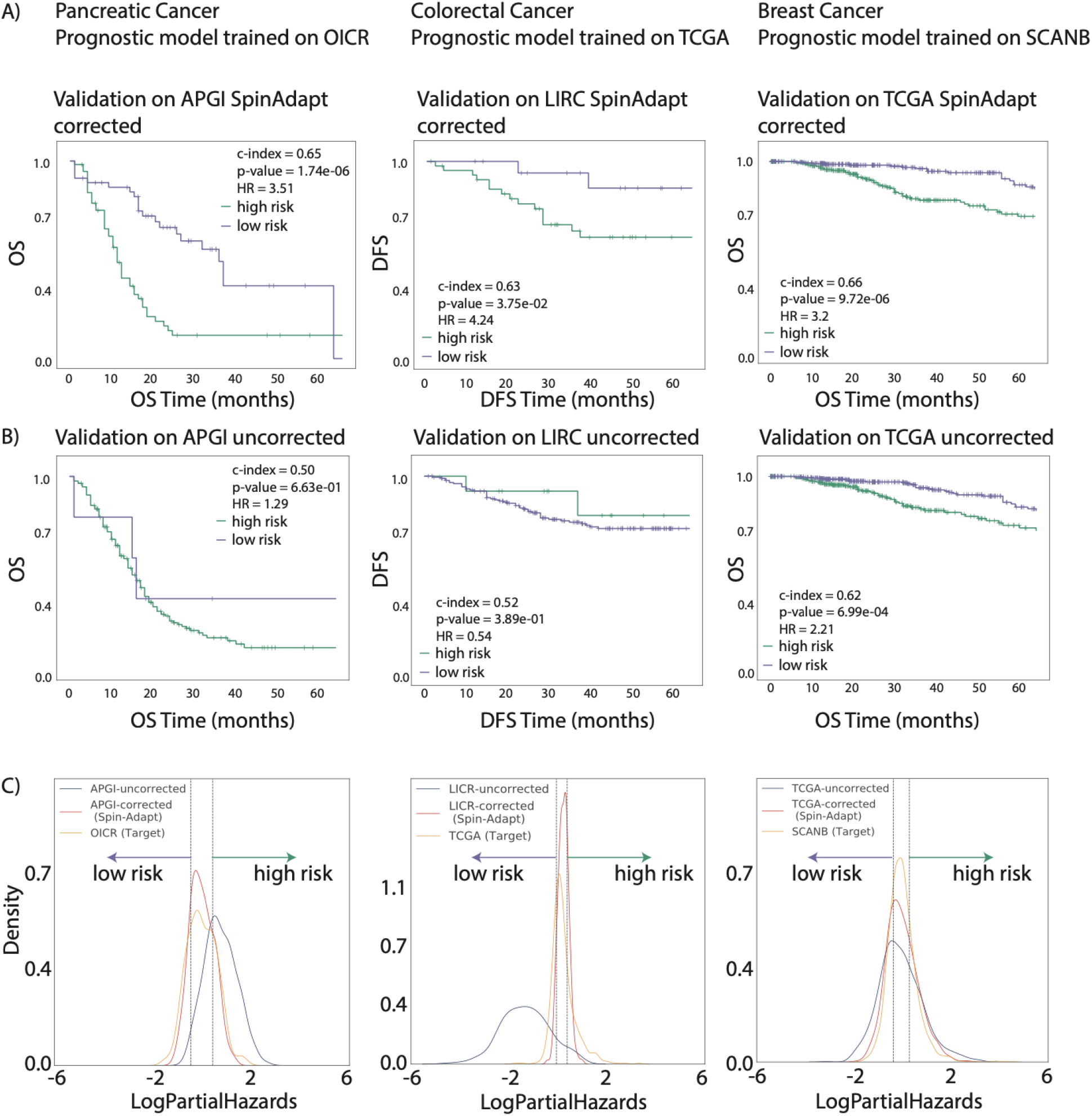
Survival curves for the predicted low-risk and predicted high-risk groups in the validation source dataset **A)** before correction and **B)** after correction using SpinAdapt, for each of the three cancer types (breast, colorectal, pancreatic). **C**) Distribution of log-partial hazards for the target-trained Cox model on the target dataset, validation source dataset, and corrected validation source dataset, for each of the three cancer types.

## Discussion

In the traditional paradigm of sharing data models across laboratories, the discussion presumes simultaneous transfer of molecular models and the associated training datasets. Therefore, all existing RNA correction methods require concurrent access to patient-level samples across all datasets. However, privacy-preserving laboratories cannot share patient-level data, and they would require a common trusted broker to have simultaneous access to their private datasets. Such trusted brokers are quite common in transactional domains such as banking where privacy and trust play a major role. Recently, the use of cryptography and distributed computing has allowed the emergence of a secure, trustless financial transaction system that eliminates the use of such brokers ^13^. Similar trust limitations still exist between healthcare organizations that actively limit data sharing due to privacy and security concerns.

SpinAdapt eliminates the need for a trusted broker, as it only operates on privacy-preserving aggregate statistics of each dataset, and allows the application of a target model on privately-held source data. Despite an inherent tradeoff between performance and privacy, SpinAdapt shows state of the art performance on diagnostic, prognostic, as well as integration tasks, outperforming similar correction algorithms that require access to private sample-level data. By only sharing data factors of the training dataset alongside the RNA-based data model, SpinAdapt allows external validation and reuse of pre-trained RNA models on novel datasets. The ability to share RNA models without the necessity of sharing model training data would improve research reproducibility across laboratories and pharmaceutical partners that cannot share patient data.

This paper demonstrates the application of SpinAdapt for transfer of diagnostic and prognostic models across distinct transcriptomic datasets, profiled across various laboratories and technological platforms, entailing RNA-Seq as well as various microarray platforms. Since the correction paradigm does not require sample-level data access, SpinAdapt enables correction of new prospective samples not included in training. The ability to correct held-out data is deemed necessary for validation frameworks, where the validation data needs to be completely held-out from training of any data models including classifiers, regressors, or batch correctors to avoid overfitting. SpinAdapt enables rigorous validation of molecular predictors across independent studies by holding-out the validation data from training of the predictor as well as the batch corrector.

In future work, the application of SpinAdapt can be empirically demonstrated in multiple OMICs data types, besides transcriptomics, for transfer of molecular predictors across technically biased datasets. The salient feature of SpinAdapt is the dependence on data factors from each dataset, which comprises the PCA factors of each dataset. Therefore, if PCA factors provide a sufficient representation of the considered OMICs dataset, SpinAdapt would be applicable for correction of technical biases across data acquisition platforms. PCA representations are common in various OMICs data modalities such as: Copy Number Variations (CNVs) ^14^, DNA Methylation ^15^, and single-cell RNA-Seq ^16^. For cases where PCA does not provide a sufficient representation of the considered data modality, PCA can be substituted with other matrix factorization methods, such as NMF, ZIFA, and pCMF, while the rest of the algorithmic details in SpinAdapt would remain unchanged.

## Acknowledgements

We thank Matthew Kase and Alexandria Bobe on the Tempus Scientific Communications team for review of the manuscript.

## Authors’ Contributions

T.A, A.A.K, E.L, M.C.S, R.P conceived the study, T.A, M.C, S.W, R.P designed and developed SpinAdapt, T.A, M.C, S.W, R.P performed computational analysis, T.A, M.C, S.W, J.R.D, A.A.S, A.A.K, M.C.S, R.P wrote the manuscript.

## Data availability

Gene expression dataset pairs (Supplementary Table 1) are compiled across various cancer types (bladder, breast, colorectal, pancreatic). Bladder cancer datasets pertain to Seiler and TCGA cohorts, where the former is downloaded from GSE87304 and the latter is obtained from ICGC under the identifier BLCA-US. Colorectal cancer datasets pertain to GSE14333 and TCGA cohorts, where the former is obtained from GSE14333 and the latter is acquired from ICGC under the identifier COAD-US. Breast cancer datasets pertain to TCGA and SCAN-B cohorts, where the former is downloaded from ICGC under the identifier BRCA-US and the latter is obtained from GSE60789. Pancreatic cancer datasets pertain to the Bailey and TCGA cohorts, which are downloaded from ICGC under the identifiers PACA-AU and PAAD-US, respectively.

## Code availability

Programming code related to data processing, algorithm design, and evaluation will be made available for non-commercial use upon request to the corresponding author.

## Methods

### Datasets

Gene expression dataset pairs are generated across various cancer types (bladder, breast, colorectal, pancreatic) pertaining to microarray platforms and RNA-sequencing (**Supplementary Table 1**). Bladder cancer datasets pertain to Seiler and TCGA cohorts ^17^, which are downloaded from GSE87304 and ICGC under the identifier BLCA-US, respectively. Colorectal cancer datasets are downloaded from GSE14333 ^18^ and ICGC under the identifier COAD-US, which are subsets of cohorts A and C, respectively, described in the ColoType prediction ^19^. Breast cancer datasets pertain to TCGA ^20^ and SCAN-B cohorts ^21^, where the former is downloaded from ICGC under the identifier BRCA-US and the latter is obtained from GSE60789. Pancreatic cancer datasets are generated using the Bailey and TCGA cohorts ^22^, which are downloaded from ICGC under the identifiers PACA-AU and PAAD-US, respectively.

The subtype labels for patients across various cancer types are generated using well-accepted subtype annotations (**Supplementary Table 1**). Bladder cancer subtypes are labeled as luminal papillary (LumP), luminal nonspecified (LumNS), luminal unstable (LumU), stroma-rich, basal/squamous (Ba/Sq), and neuroendocrine-like (NE-like), generated using a consensus subtyping approach ^17^. Breast cancer subtypes are labeled as Luminal A (LumA), Luminal B (LumB), HER2-enriched (Her2), and Basal-like (Basal) ^23^. Colorectal cancer subtypes are labeled as CMS1, CMS2, CMS3, CMS4, as published by Colorectal Cancer Subtyping Consortium (CRCSC) ^24^. Pancreatic cancer subtypes are generated using expression analysis, labeled as squamous, pancreatic progenitor, immunogenic, and aberrantly differentiated endocrine exocrine (ADEX) ^25^.

### Preprocessing

For each cancer type, we only keep patients with both expression data and subtype annotation labels available. Molecular subtypes with less than 5 patients in any cancer cohort are removed. For the microarray expression datasets (GSE14333 and Seiler, see **Supplementary Table 1**), multiple probe sets may map to the same gene. The expression values were averaged across such probe sets to get gene expression value. Furthermore, for each cancer type, we remove genes with zero variance, and we only keep genes common between source and target, sorted in alphabetical order. Finally, we normalized the RNA-Seq datasets using the variance stabilizing transform (VST) from DeSeq2, whereas microarray data were not normalized beyond their publication.

### Benchmarking methods

For Seurat, we used the default package parameters, except when the number of samples in either dataset is less than 200, where the default value of k.filter does not work. Therefore, when the number of samples in either dataset is less than 200, we set the k.filter parameter to 50. For ComBat, a design matrix is created using the batch labels, and the method is implemented using the sva package version 3.34.0. Similarly, Limma is implemented using limma package version 3.42.2. For Seurat and Scanorama, we deployed package versions 3.2.2 and 1.6, respectively.

### Parameters for SpinAdapt

For any given pair of source and target datasets, let *p* be the number of genes, *n*_*s*_ be the number of samples in source, and *n*_*t*_ be the number of samples in target. Across all experiments performed in this study, the parameters in SpinAdapt (Algorithm 1) are set as follows: α = 0. 01, λ = (2/3) * *min*(*n*_*s*_, *n*_*t*_), and *variance*_*norm*_ is set to True when the source is microarray but False otherwise. However, for integration analysis of source and target datasets, we always set *variance*_*norm*_ = *True*.

### Evaluation methods for transfer of diagnostic models

For each of the 17 cancer subtypes (see **Supplementary Table 1**), we train a one-vs-rest random forest classifier on the target dataset, such that the classifier learns to discriminate the selected subtype against all other subtypes in the target. Specifically, all target samples annotated with the selected subtype are given a positive label, while the rest of the target samples are assigned a negative label. The hyperparameters for the random forest classifier are learnt in a three-fold cross-validation experiment on the target dataset.

We compare multiple batch correction (adaptation) methods for transfer of the subtype classifier from target to the source dataset. The transfer requires adaptation of the source dataset to the target reference. For unbiased performance evaluation of a batch correction method, the test set for the classifier and the training set for the correction method need to be disjoint, and thus, the correction model does not train on the classifier test set. We propose a framework for validating transfer of classifiers across datasets that avoids such information leakage.

The validation framework randomly splits the source dataset into two mutually-exclusive subsets: source-A and source-B. First, the adaptation model is trained from source-A to target fit), and applied to source-B (transform). The target classifier generates predictions on corrected source-B (**Supplementary Figure 1A**). Second, the adaptation model is fit from source-B to target, followed by transformation of source-A and generation of predictions on corrected source-A (**Supplementary Figure 1B**). Finally, the classification performance is quantified by computing F-1 scores for all samples in the held-out corrected source-A and source-B subsets. We repeat the entire procedure 30 times, choosing a different partitioning of source into source-A and source-B, and we report the mean F-1 score for each subtype over the 30 iterations.

We evaluate SpinAdapt using the aforementioned framework for validating transfer of subtype classifiers. However, since Seurat and ComBat cannot transform out-of-sample data, they can only correct samples included in training of these correction methods. Therefore, for these two methods, we train the correction model on the classifier test set, followed by application of target-trained classifier on the corrected test set. Specifically, for ComBat and Seurat, we fit-transformed source-A to target, fit-transformed source-B to target, and computed F-1 scores for all samples in the transformed source-A and source-B subsets. As before, we repeat the procedure 30 times, using the same data splits as used for SpinAdapt validation, which enabled pairwise performance comparisons between SpinAdapt, Seurat, and ComBat.

For each subtype, we performed the two-sided paired McNemar test to identify if the differences between any pair of adaptation methods are statistically significant ^26^. Due to the rarity of positives for a selected subtype in each dataset, we perform the McNemar test only on samples with positive ground truth. Each positive sample is assigned a correct or incorrect classification label. Then, for each pair of correction methods, the McNemar test statistic is evaluated on the disagreements between correction methods on the positive samples. We report the median *P*-value across the 30 repetitions of the validation framework (**Supplementary Table 4**).

### Evaluation methods for dataset integration

A common task for RNA-based algorithms is dataset integration (batch mixing). There is an inherent trade-off between batch mixing and preservation of the biological signal within integrated datasets. To quantify preservation of the biological signal, we quantify subtype-wise separability (no mixing of tumor subtypes) in the integrated datasets. Therefore, for high data integration performance, we want to minimize subtype mixing while maximizing batch mixing.

To compare SpinAdapt with other batch integration methods, we assess the goodness of batch mixing and tissue type separation. First, we employed the average silhouette width (ASW) to quantify batch mixing and tissue segregation. The silhouette score of a sample is obtained by subtracting the average distance to samples with the same tissue label from the average distance to samples in the nearest cluster w.r.t. the tissue label, and then dividing by the larger of the two values ^27^. Therefore, the silhouette score for a given sample varies between -1 and 1, such that a higher score implies a good fit among samples with the same tissue label, and vice versa. In other words, a higher average silhouette width implies mixing of batches within each tissue type or/and separation of samples from distinct tissue types.

To explicitly quantify batch mixing and tissue segregation, independently, we employ the local inverse Simpson’s index (LISI). The LISI metric assigns a diversity score to each sample by computing the effective number of label types in the local neighborhood of the sample. Therefore, the notion of diversity depends on the label under consideration. When the label is set to batch membership, the resulting metric is referred to as batch LISI (bLISI), since it measures batch diversity in the neighborhood of each sample. When the label is set to tissue type, the resulting metric is referred to as tissue LISI (tLISI), since it measures tissue type diversity in sample neighborhood. For good integration, we sought sample neighborhoods with high batch diversity and low subtype diversity, which correlates with high bLISI and low tLISI score, respectively. For each integration method and cancer dataset, we report average bLISI and tLISI scores across all samples in source and target datasets (**Supplementary Table 3, Supplementary Figure 4)**. When comparing methods using average bLISI, which measures dataset mixing, Seurat outperforms SpinAdapt on breast and bladder cancer datasets (*P* < 10e-3), whereas SpinAdapt outperforms ComBat, Limma, and Scanorama on colorectal and pancreatic cancer datasets (*P* < 10e-3) (**Supplementary Tables 3 and 5**). When comparing methods using average tLISI, SpinAdapt significantly outperforms all other methods on breast (*P* < 10e-13), colorectal (*P* < 10e-7), and pancreatic (*P* < 0.05) cancer datasets (**Supplementary Tables 3 and 5**), implying SpinAdapt best preserves molecular structures for dataset integration. When comparing methods using tLISI on the bladder cancer dataset, SpinAdapt outperforms Seurat, ComBat, Limma (P < 10e-6), whereas Scanorama outperforms SpinAdapt without significance.

The various integration metrics including silhouette, bLISI, and tLISI scores are computed on the UMAP embeddings of the integrated datasets for each cancer type (**Supplementary Table 1**). Specifically, the scores in each experiment are computed on the first 50 components of the UMAP transform, where the UMAP embeddings are computed using default parameters of the package. The average silhouette width, bLISI, and tLISI scores are reported along with the standard errors (**Supplementary Table 3**). For each metric, significance testing between methods is performed by a two-sided paired Wilcoxon test (**Supplementary Table 5**).

### Evaluation methods for transfer of prognostic models

For each of the four cancer datasets (Colorectal, Breast, Pancreatic, see **Supplementary Table 6**), we trained a Cox Proportional Hazards model on the target dataset using a gene signature determined through an ensemble method performed on the target dataset. The ensemble method for feature selection uses four ranked lists of genes, based on different statistical tests or machine learning models: Chi-square scores, F-scores, Random Forest importance metrics, and univariate Cox PH p-value. We tested the predictive values of various permutations of genes of increasing length (n = 10, 50, 75, 100, 200, 300, 500 genes) as signatures of a Cox PH model trained and tested on random splits of the target dataset, in a five-fold cross-validation setting, where 50% of the target dataset was assigned to the training set and the remainder was assigned to the test set. The best performing signature was determined based on the c-index determined on the five random test sets. We then used this signature to train a final Cox PH model on the target dataset.

### Visualization

We employ the UMAP transform to visualize the batch integration results for each cancer type (**Figures 2 and 4, Supplementary Figure 3**). Specifically, we perform visualization in each experiment using the first two components of the UMAP embeddings, where the number of neighbors are set to 10 and the min_dist parameter is set to 0.5. These parameters are fixed for all visualizations in the study that employ UMAP embeddings.

### Algorithm Details

SpinAdapt inputs source and target expression datasets for training, corrects the batch-biased source expression data, even when the source data is held-out from training, followed by evaluation of the target-trained predictor on the corrected source data. The algorithm, as outlined in Algorithm 1, can be broken down into several main steps: computation of source and target data factors from source and target datasets in Steps 1 and 2, respectively, estimation of a low-rank affine map between source and target PCA basis in Step 3, adaptation (correction) of the source dataset in Step 4, and finally, evaluation of the target-trained predictor on adapted source dataset in Step 5. Notably, Step 4, Algorithm 1 can adapt source dataset *X*_*sh*_ that is held-out from the training source dataset *X*_*s*_. Steps 1 and 2 are executed using Algorithm 2, whereas Steps 3 and 4 are executed using Algorithms 3 and 4, respectively. These steps are explained next in further detail.

In Step 1, Algorithm 1, data factors are computed for the source dataset *X*_*s*_, where the data factors comprise of the PCA basis *U*_*s*_, gene-wise means *m*_*s*_, and gene-wise variances *s*_*s*_ of the source dataset. The details for the computation of these data factors are outlined in Algorithm 2, where the gene-wise means and variances are computed in Steps 2-3, whereas the PCA basis are computed in Steps 4-6. Similarly, data factors are computed for the target dataset in Step 2, Algorithm 1, where the data factors entail the PCA basis *U*_*t*_, gene-wise means *m*_*t*_, and gene-wise variances *s* for the target dataset *X*_*t*_. The gene-wise means and variances *m*_*s*_, *m*_*t*_, *s*_*s*_, *s*_*t*_ are used in the correction step (Step 4, Algorithm 1), whereas the PCA basis *U*_*s*_ and *U*_*t*_ are used in the train and correction steps (Steps 3-4, Algorithm 1). The usage of statistics *s*_*s*_ and *s*_*t*_ in Step 4, Algorithm 1 is optional depending on the boolean value of *variance*_*norm*_, as we explain later.

Notably, Algorithm 1 does not require simultaneous access to sample-level patient data in source and target datasets at any step. Computation of source data factors in Step 1 needs access to *X*_*s*_ only, whereas computation of target data factors in Step 2 needs access to *X*_*t*_ only. Training in Step 3 only requires access to the PCA basis of source and target datasets. Since the PCA basis cannot be used for recovery of sample-level patient data, the basis are privacy-preserving (**Supplementary Note**). Adaptation in Step 4 requires access to the source expression data *X*_*s*_, linear map *A*, and the data factors of both datasets, without requiring access to the target dataset *X*_*t*_.

#### Algorithm 1: SpinAdapt algorithm

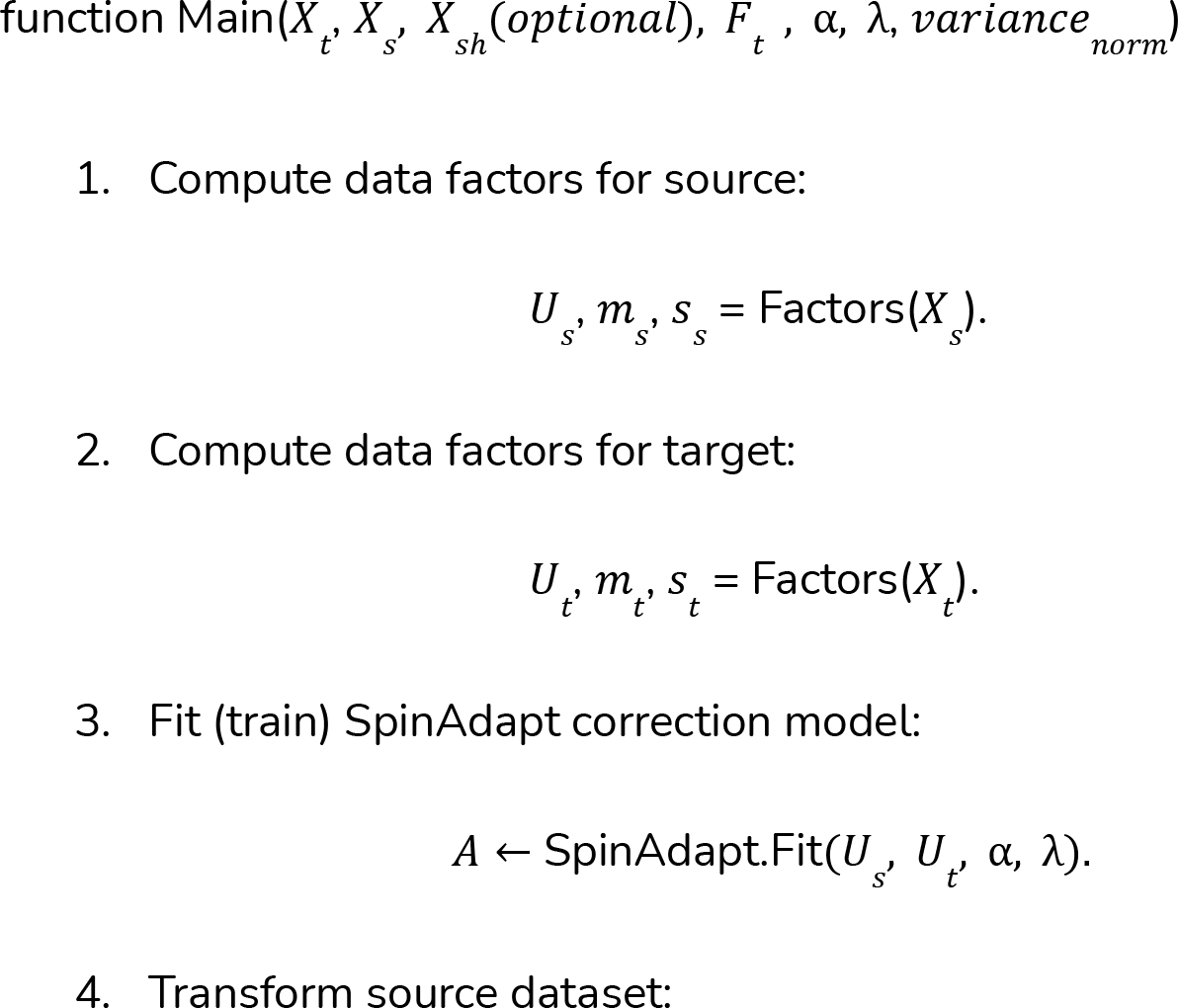
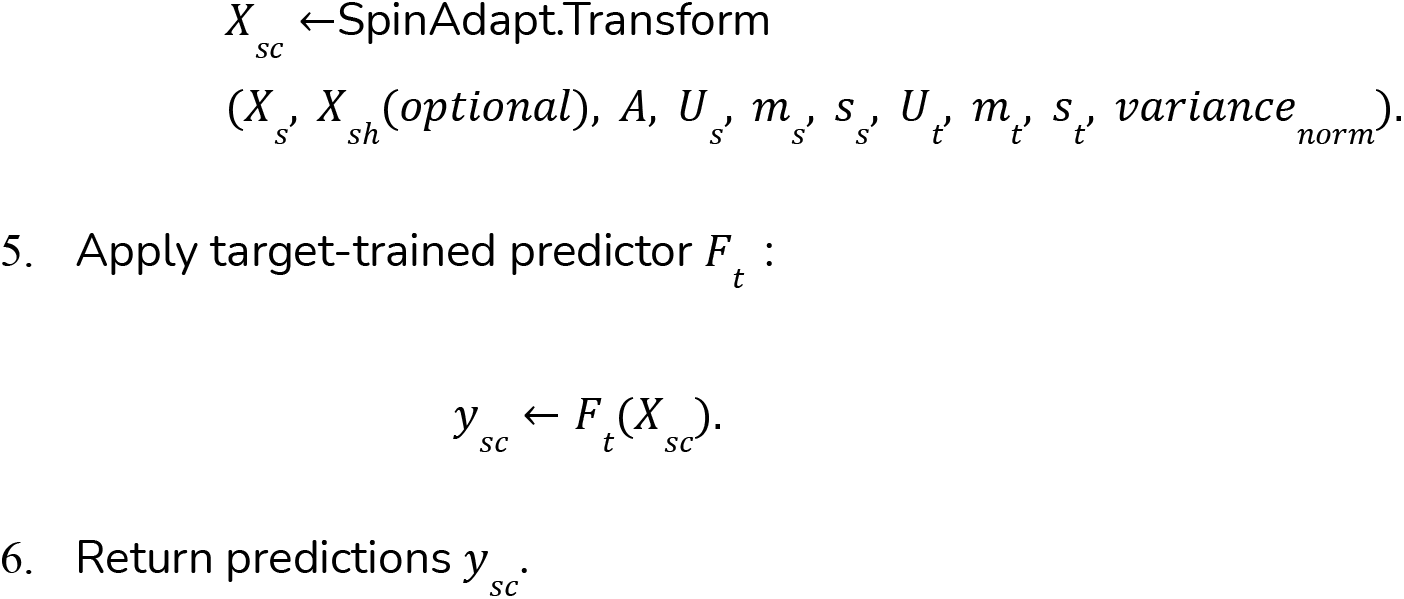

#### Algorithm 2: Compute data factors, gene-wise means and variances

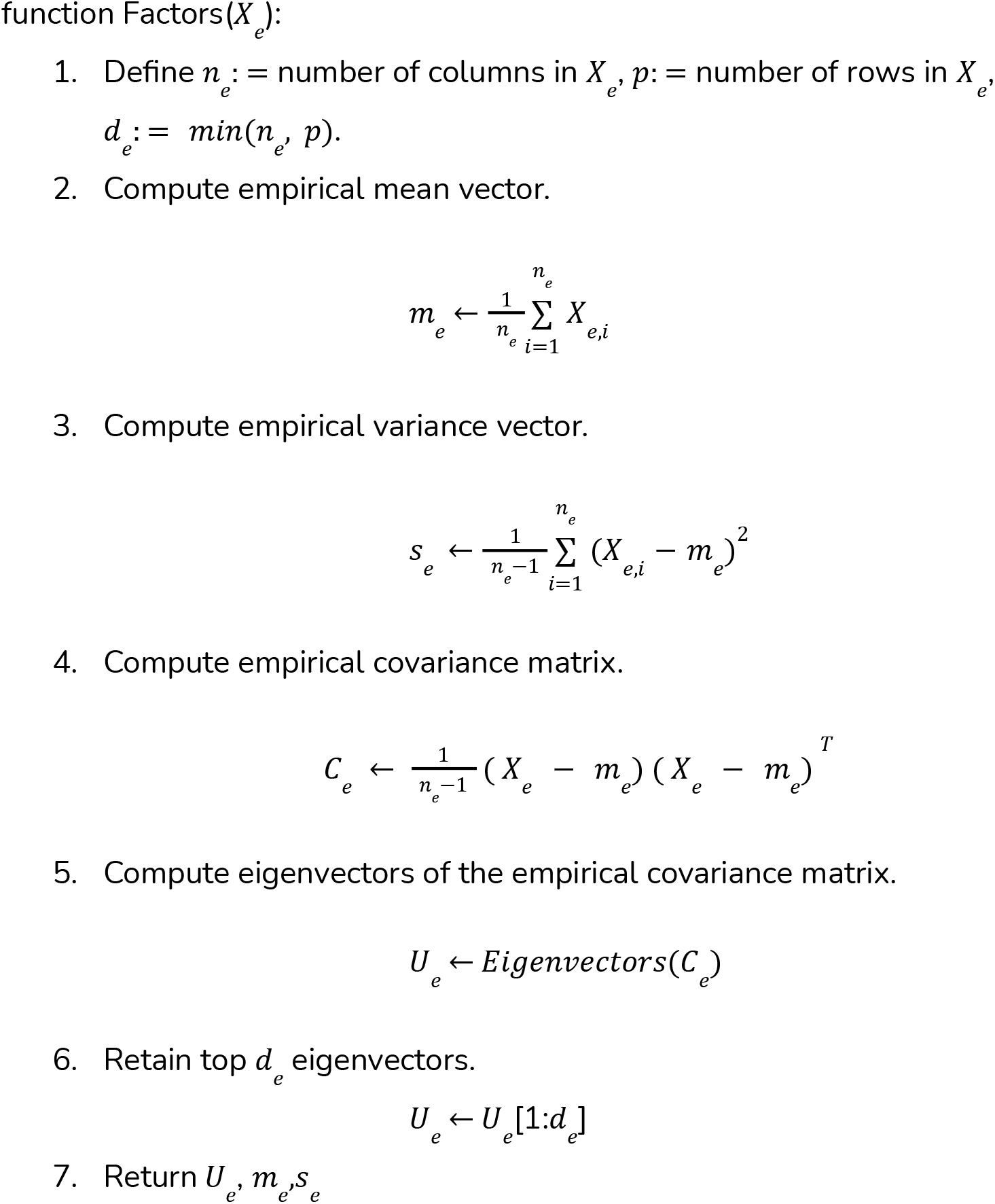

#### Algorithm Details: Glossary

We define the data structures employed in Algorithm 1, with the dimensionality of each structure. The dimensionality is stated in terms of *p*, the number of genes; *n*_*s*_, number of samples in source dataset; *n*_*t*_, number of samples in target dataset; *d*_*s*_, dimensionality of source latent space; *d*_*t*_, dimensionality of source latent space.

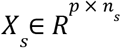: The train source dataset
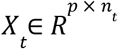: The train target dataset
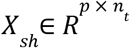: The held-out source dataset
*X*_*s,i*_ ∈ *R*^*p*^: The i-th column of *X*_*s*_
*X*_*t, i*_ ∈ *R*^*p*^: The i-th column of *X*_*t*_
*m*_*s*_∈ *R*^*p*^: The empirical gene-wise mean of source dataset
*m*_*t*_∈ *R* ^*p*^: The empirical gene-wise mean of target dataset
*s*_*s*_∈ *R* ^*p*^: The empirical gene-wise variance of source dataset
*s*_*t*_∈ *R* ^*p*^: The empirical gene-wise variance of target dataset
*C*_*s*_∈ *R* ^*p* × *d*^: The empirical covariance of source dataset
*C*_*t*_∈ *R* ^*p* × *d*^: The empirical covariance of target dataset
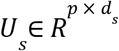: Principal Component factors for source dataset
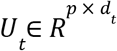: Principal Component factors for target dataset
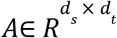: Transformation matrix
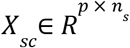: The corrected output source dataset
*X* (*i,j*): The i-th row and j-th column of any matrix *X*
*ν*(*i*): The i-th entry of any vector *ν*
*F*_*t*_: Classifier trained on the target dataset

The parameters α and λ correspond to step size and regularization parameters for the iterative algorithm Fit (Algorithm 3). The parameter *variance*_*norm*_ is a boolean variable, which determines if the adaptation step (Step 4, Algorithm 1) entails variance-normalization of the source dataset (see Algorithm 4 for details).

### Algorithm Details: Learning transformation between PCA factors

In Step 3, Algorithm 1, we learn a low matrix-rank transformation between PCA factors of the source dataset and PCA factors of the target dataset. We pose a non-convex optimization problem to learn the transformation, and then we present an effective computational approach to solve it, as we explain next.

#### Objective function for Step 3, Algorithm 1

The objective function is based on Frobenius norm between transformed source PCA basis *U*_*s*_*A* and the target PCA basis *U*_*t*_, as follows

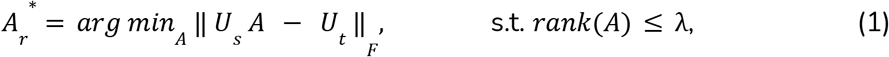

where A represents the transformation matrix, λ represents the matrix-rank constraint, and rank(A) represents the matrix-rank of A. In the main term, it can be seen that the *i*-th column of the transformation matrix A determines what linear combination of the columns of *U*_*s*_ best approximates the *i*-th column of *U*_*t*_, where *i* = 1, 2,…, *d*_*t*_. Therefore, the intuition behind the main term is to approximate each target factor using some linear combination of source factors.

The inequality constraint in equation (1) is a matrix-rank penalization term, which restricts the solution space of A to matrices with matrix-rank less than λ. The rank constraint is reminiscent of sparse constraint in sparse recovery problems, where the constraint restricts the maximum number of non-zero entries in the estimated solution, thereby reducing the sample complexity of the learning task. Similarly, in equation (1), the constraint restricts the maximum matrix-rank of A, making the algorithm less prone to overfitting, while decreasing the sample requirement of learning the affine map from source to target factors. However, the problem posed in equation (1) turns out to be non-convex, and thus hard to solve. We employ traditional optimization techniques and derive an efficient routine for computing *A*_*r*_^*^, as follows in the next subsection.

#### Optimization solution for Step 3, Algorithm 1

Let g(A) = ‖ *U*_*s*_ *A* − *U*_*t*_ ‖_*F*_, and let 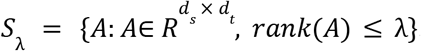. Then, the objective function in equation (1) can be re-written as

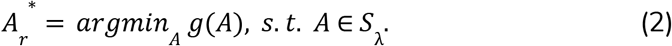

Gradient descent can be used to minimize *g*(*A*) w.r.t. *A* because the function is convex and differentiable. In contrast, equation (2) cannot be evaluated using gradient descent, since the set of low-rank matrices *S*_λ_ is non-convex. However, we note that the Euclidean projection onto the set *S*_λ_ can be efficiently computed, which hints that equation (2) can be minimized using projected gradient descent, as we explain next. Let the Euclidean projection of a matrix *A* onto set *S*_λ_ be denoted by *P*_λ_ (*A*). Then, mathematically we have *P*_λ_ (*A*) = *argmin*_*Z*_ {‖ *A* − *Z* ‖_*F*_ : *Z* ∈ *S*_λ_}. From the Eckart-Young Theorem, we know that *P*_λ_(*A*) can be efficiently evaluated by computing the top λ singular values and singular vectors of *A*. The closed-form solution of *P*_λ_ (*A*) is given by the SVD transform *U*_λ_ Σ_λ_ *V*_λ_ ^*T*^, where columns of *U*_λ_ contain the top λ eigenvectors of *AA*^*T*^columns *V*_λ_ of contain the top λ eigenvectors of *A*^*T*^*A*, and entries of the diagonal matrix Σ_λ_ are square roots of the top λ eigenvalues of *AA*^*T*^. We are finally ready to present an algorithm for solving (2).

#### Pseudocode for SpinAdapt.Fit (Algorithm 3)

We present the algorithm for executing Step 3, Algorithm 1, which is essentially a solution to the optimization problem in (2). We propose the use of the projected gradient descent algorithm to evaluate (2), which is an iterative application of the following descent step:

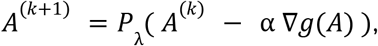

for *k* = 0, 1, 2, 3, … till convergence. Details are provided in Algorithm 3 below.

##### Algorithm 3 Learn transformation from source to target factors (Fit)

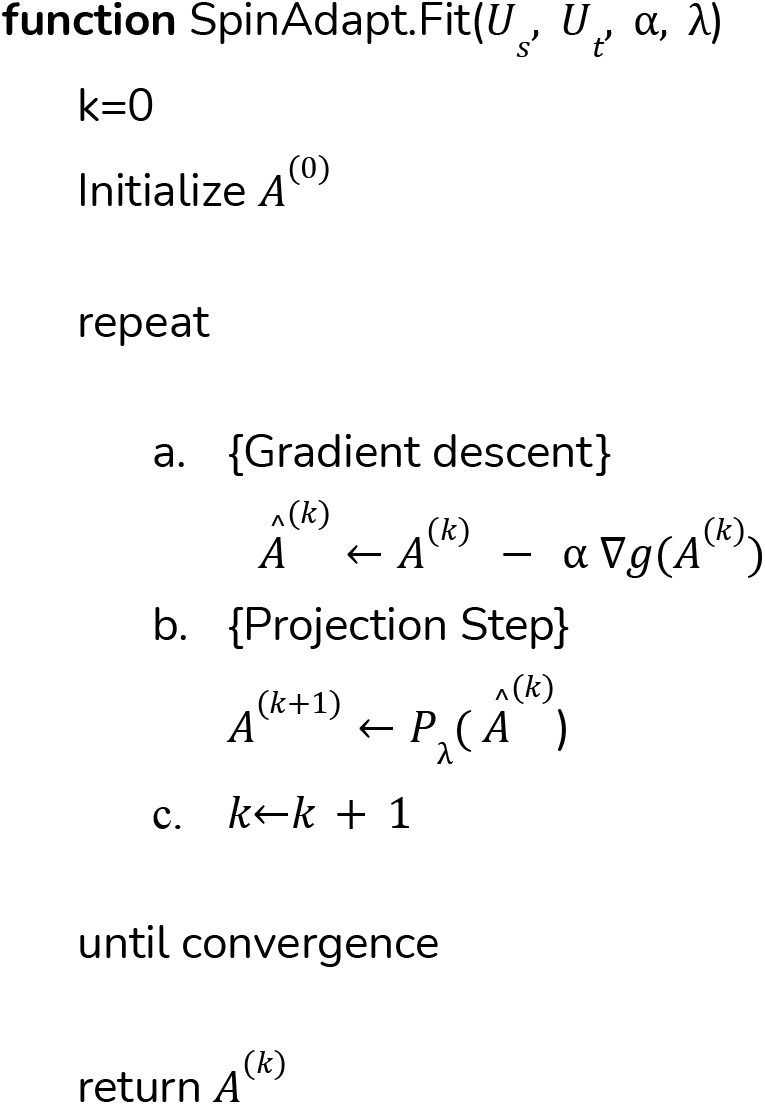

#### Pseudocode for SpinAdapt.Transform (Algorithm 4)

Finally, we present the algorithm for executing Step 4, Algorithm 1, where the batch-biased dataset is corrected using the transformation *A*. Details are outlined in Algorithm 4, as follows. In Step 1, Algorithm 4, the held-out source dataset *X*_*sh*_ is selected for correction, if provided. If held-out evaluation data (*X*_*sh*_) is not provided, the train source dataset *X*_*s*_ is selected for correction. In Step 2, if the input parameter *variance*_*norm*_is set to True, the variance of each gene in the source dataset is matched to variance of the corresponding gene in the target dataset. In Step 3, the PCA embeddings of each source sample are computed. In Step 4, the computed PCA embeddings are corrected, using the transformation matrix *A*. In Step 5, the corrected PCA embeddings are transformed to the gene expression space. Finally, the corrected source gene expression profiles are returned in Step 6.

##### Algorithm 4 Adapt the source dataset (Transform)

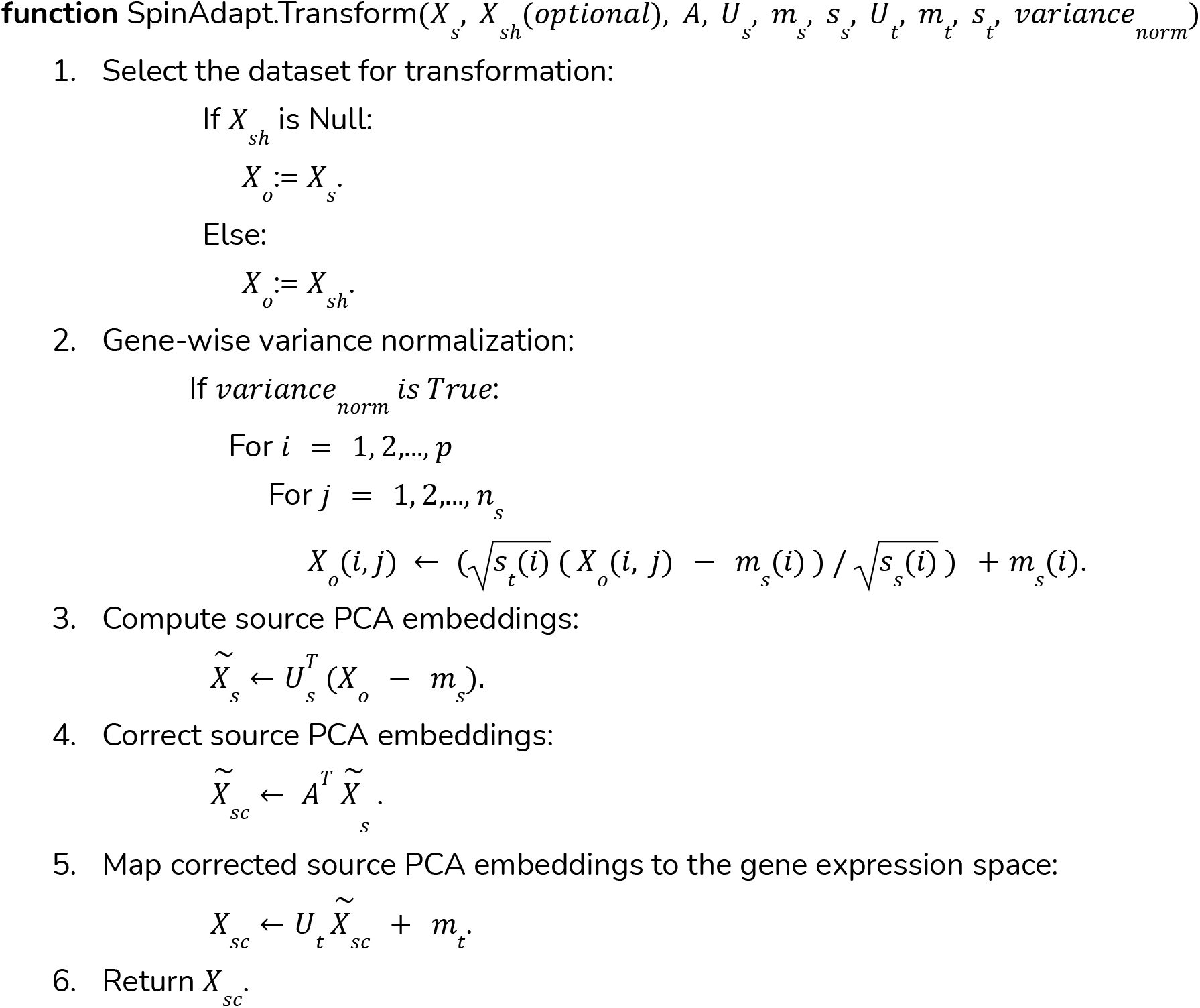

