## Supplementary Files for "Privacy Preserving RNA-Model Validation Across Laboratories"

### Supplementary figures

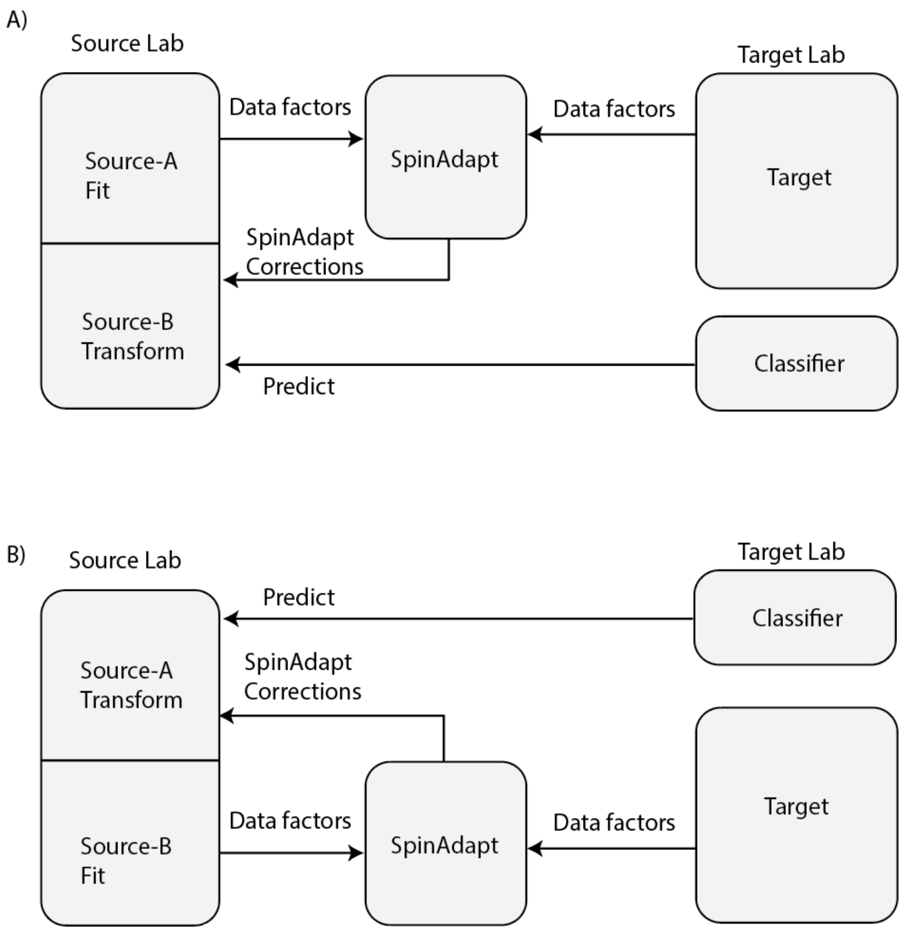


**Supplementary Figure 1.** Experimental framework for the validation of the target-trained molecular predictors on batch-corrected source dataset, such that the correction model never trains on the test data for the predictor.

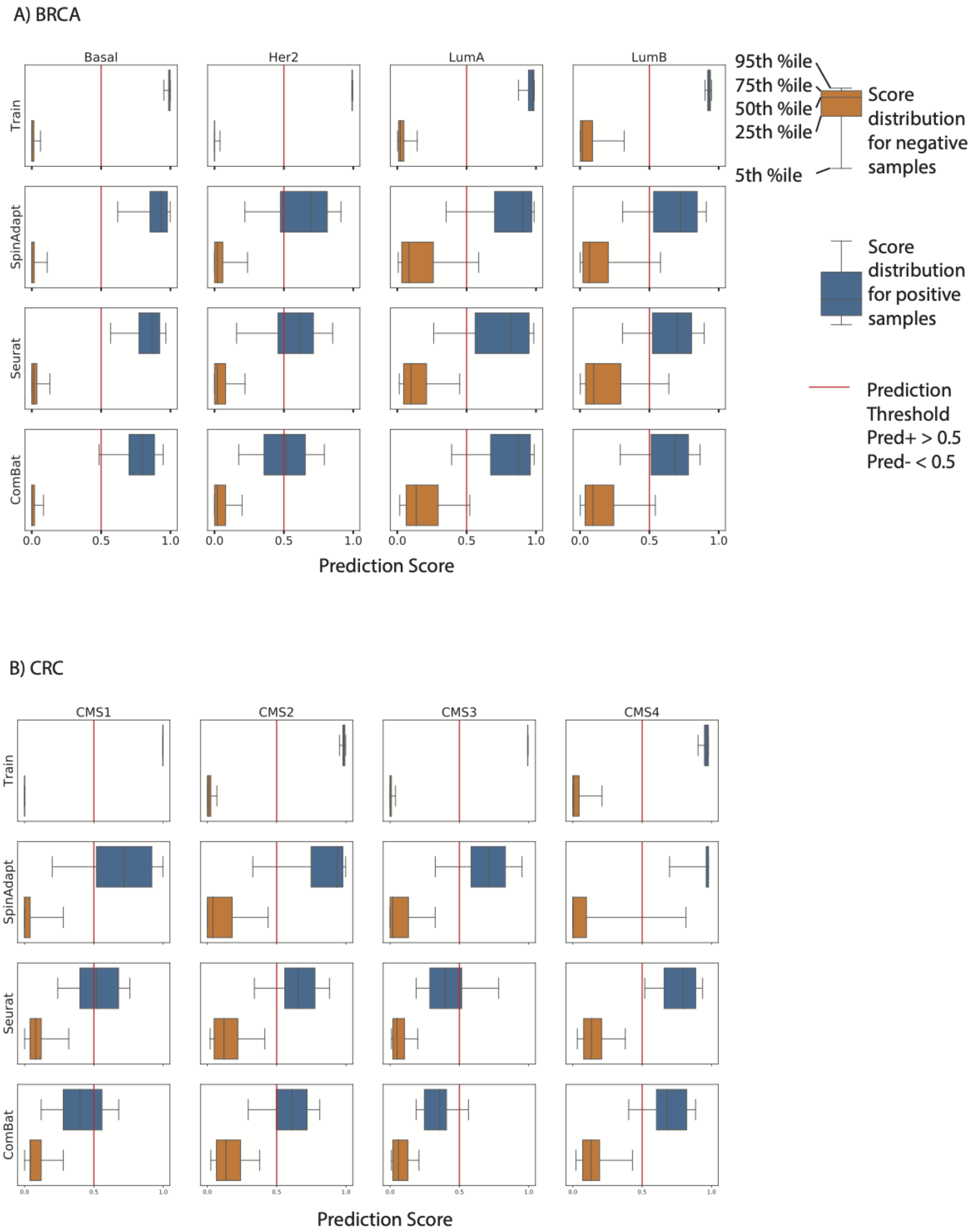
A) BRCA
Basal
Her2
LumA
LumB
Train
SpinAdapt
Seurat
ComBat
0.0
0.5
1.0
Prediction Score
95th %ile
75th %ile
50th %ile
25th %ile
5th %ile
Score distribution for negative samples
Score distribution for positive samples
Prediction Threshold
Pred+ > 0.5
Pred- < 0.5
B) CRC
CMS1
CMS2
CMS3
CMS4
Train
SpinAdapt
Seurat
ComBat
0.0
0.5
1.0
Prediction Score

C) PANC

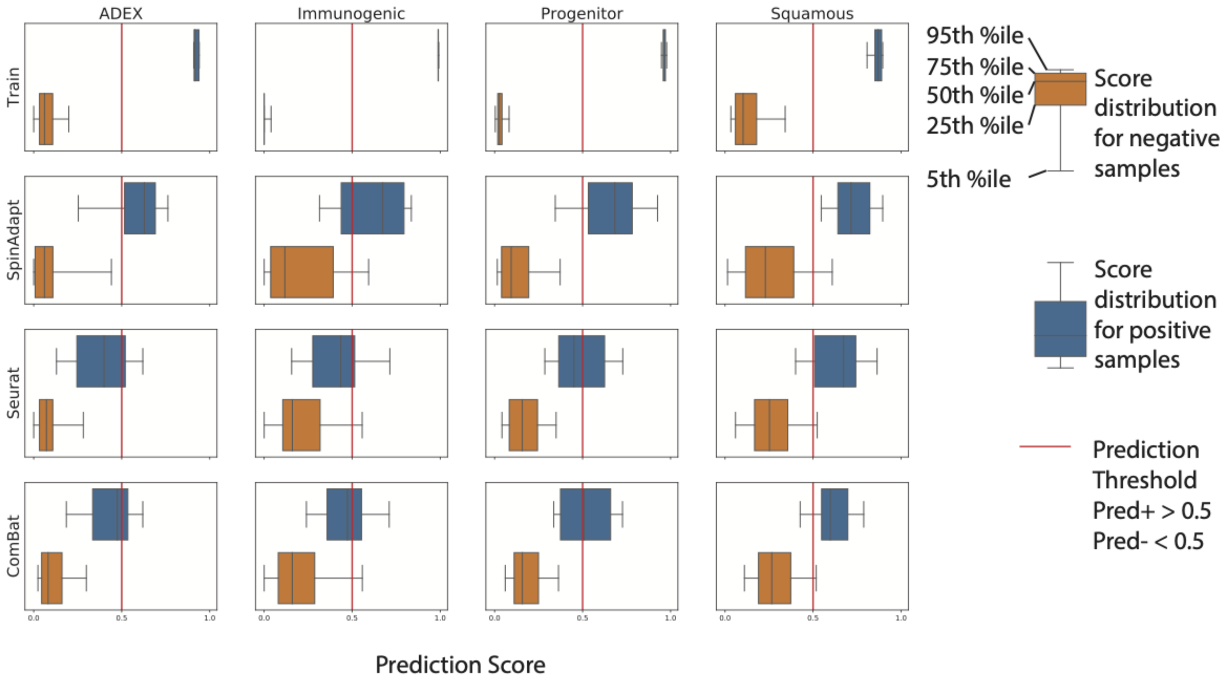


D) Bladder

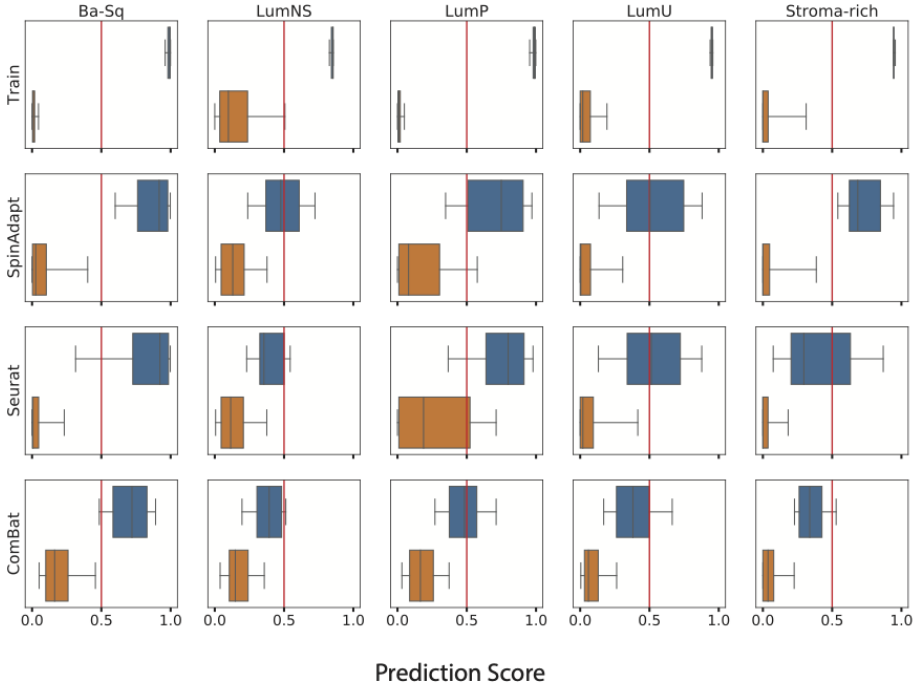


**Supplementary Figure 2.** Boxplots for predictor scores on batch-corrected source dataset for each cancer subtype. For each cancer subtype (column) and correction method (row), the boxplots for positive and control samples are plotted separately, and the vertical line represents the decision threshold at 0.5 (**methods**). Better performance is achieved when control and positive sample score distributions are shifted to the left and right, respectively. **A**) Breast: SpinAdapt obtains higher median test scores on the positive samples, **B**) Colorectal: the SpinAdapt test score distributions for positive samples are shifted to the right, but CMS4 subtype suffers from lower specificity, **C**) Pancreatic: the SpinAdapt test score distributions for positive samples are shifted to the right, compared with Seurat and ComBat. **D**) Bladder: SpinAdapt obtains lower median test scores on the control samples for LumP subtype, while obtaining higher median test scores on the positive samples for the LumNS and Stroma-rich subtypes, compared with Seurat and ComBat.

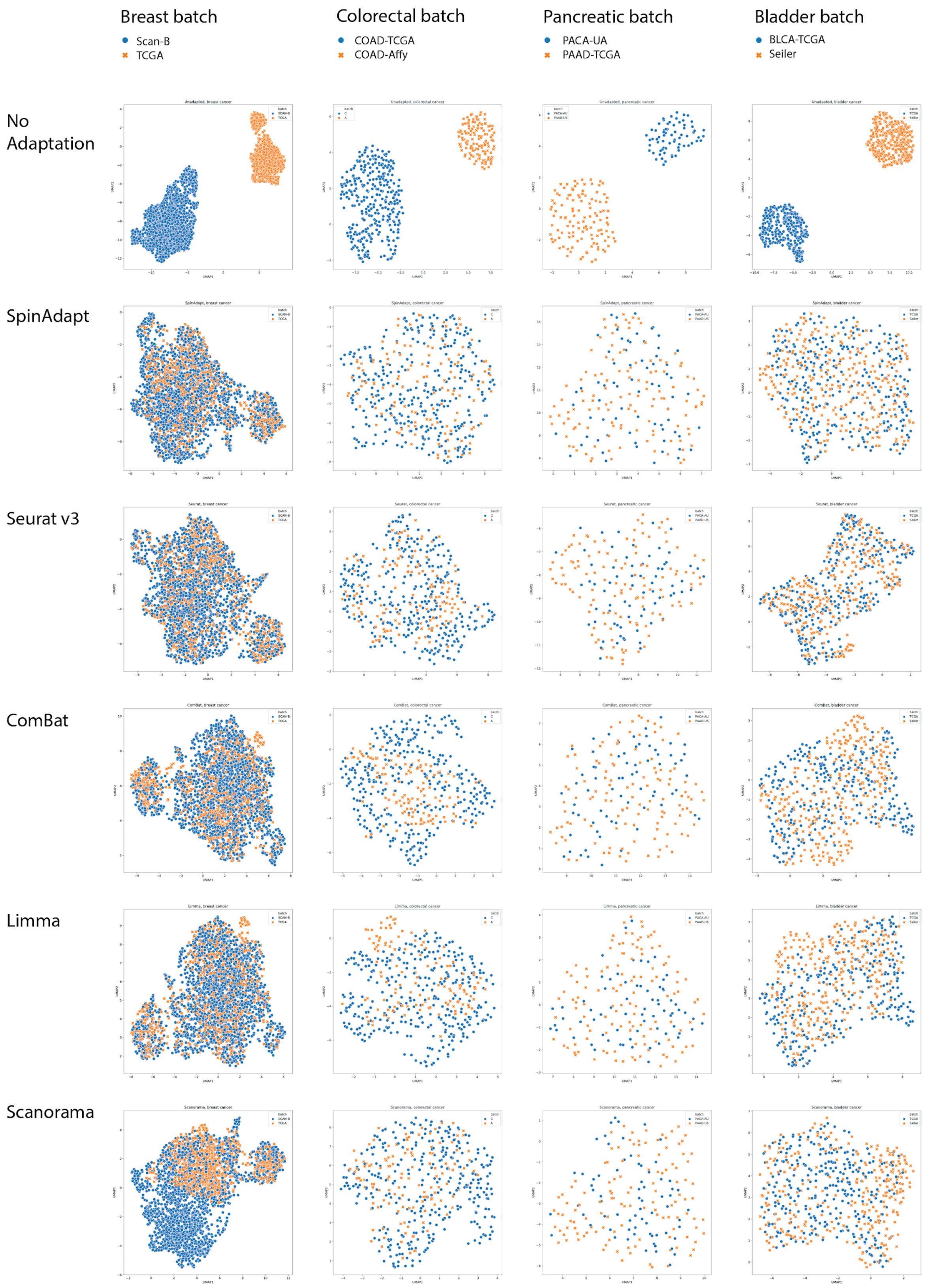


**Supplementary Figure 3. A**) UMAP plots for dataset integration, labeling samples by dataset.

Top panel shows dataset-based clustering in each cancer dataset before integration. Integration requires good batch mixing within integrated datasets, which is achieved by most methods. ComBat, Limma, Scanorama are outperformed by Seurat, SpinAdapt in terms of batch mixing in Colorectal, while Scanorama achieves poor batch mixing in Breast.

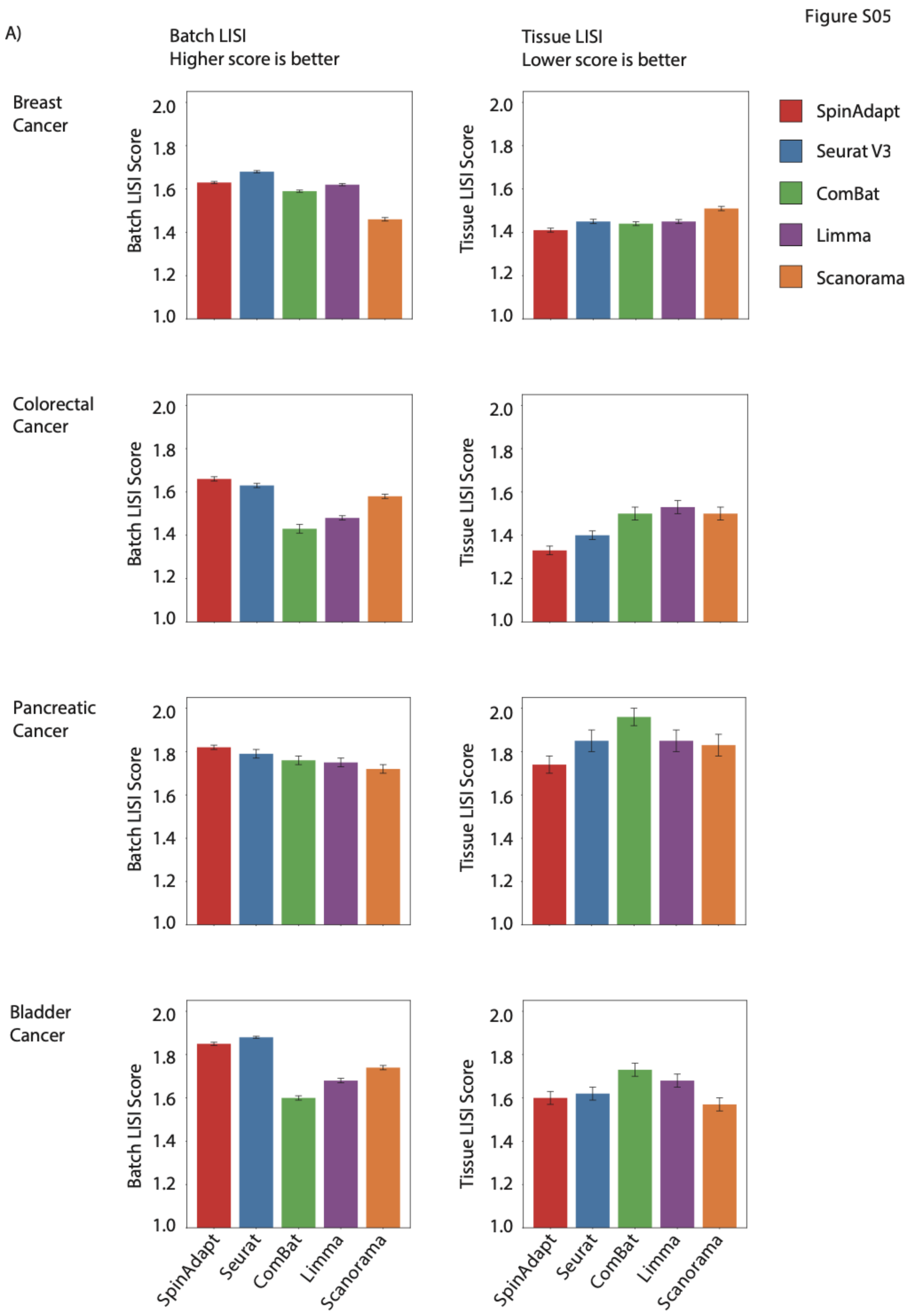


**Supplementary Figure 4.** A) Quantification of dataset integration performance using batch LISI (bLISI) and tissue LISI (tLISI) metrics, where bLISI measures batch homogeneity (a higher score is better) and tLISI measures subtype heterogeneity in local sample neighborhoods (a lower score is better). For each dataset, correction method, and performance metric, the associated barplot reports the mean and standard error over all the integrated samples.

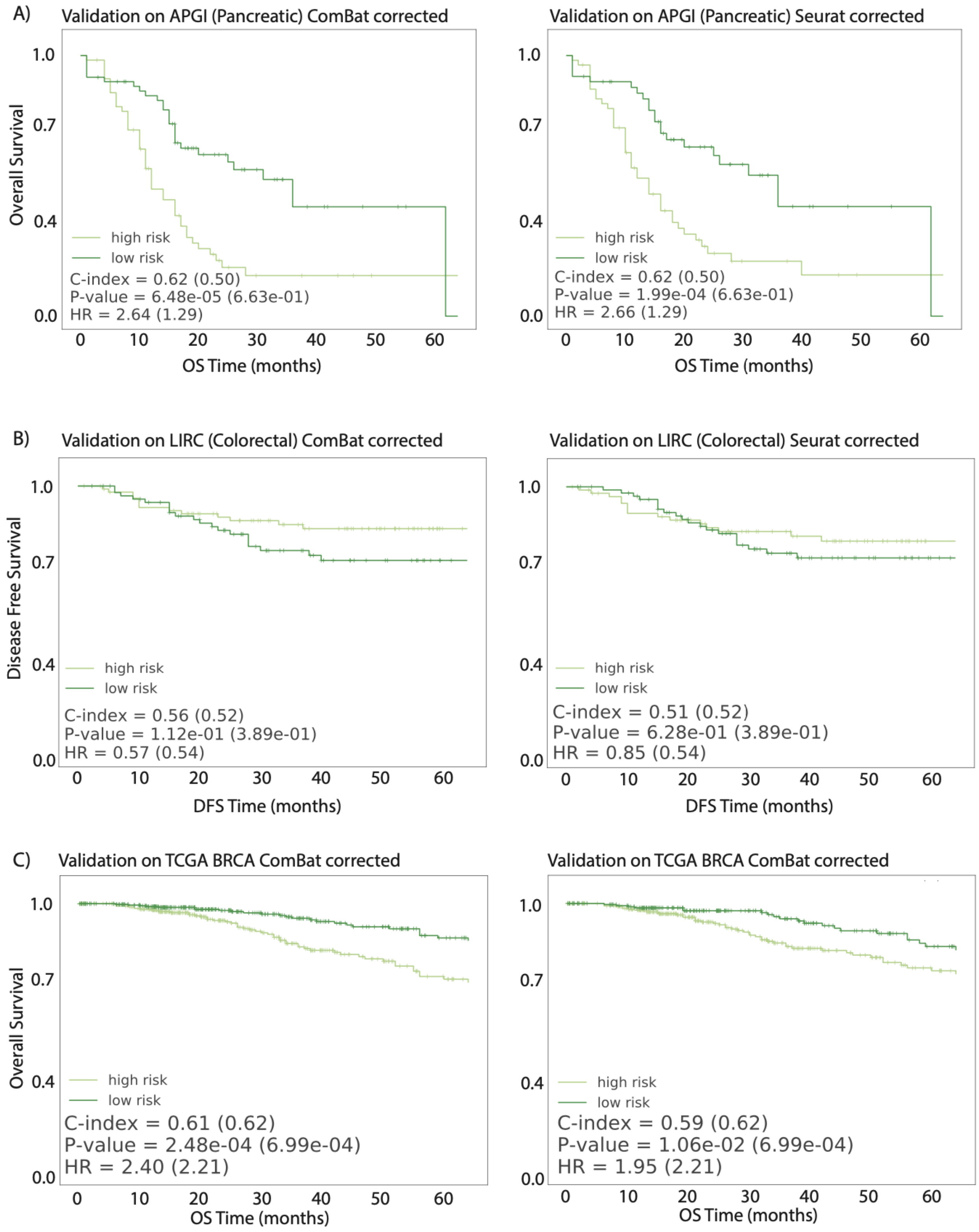


**Supplementary Figure 5.** Survival curves for predicted low-risk and high-risk groups on the validation (source) dataset for **A)** pancreatic, **B)** colorectal, and **C)** breast cancer types, using Seurat and ComBat.

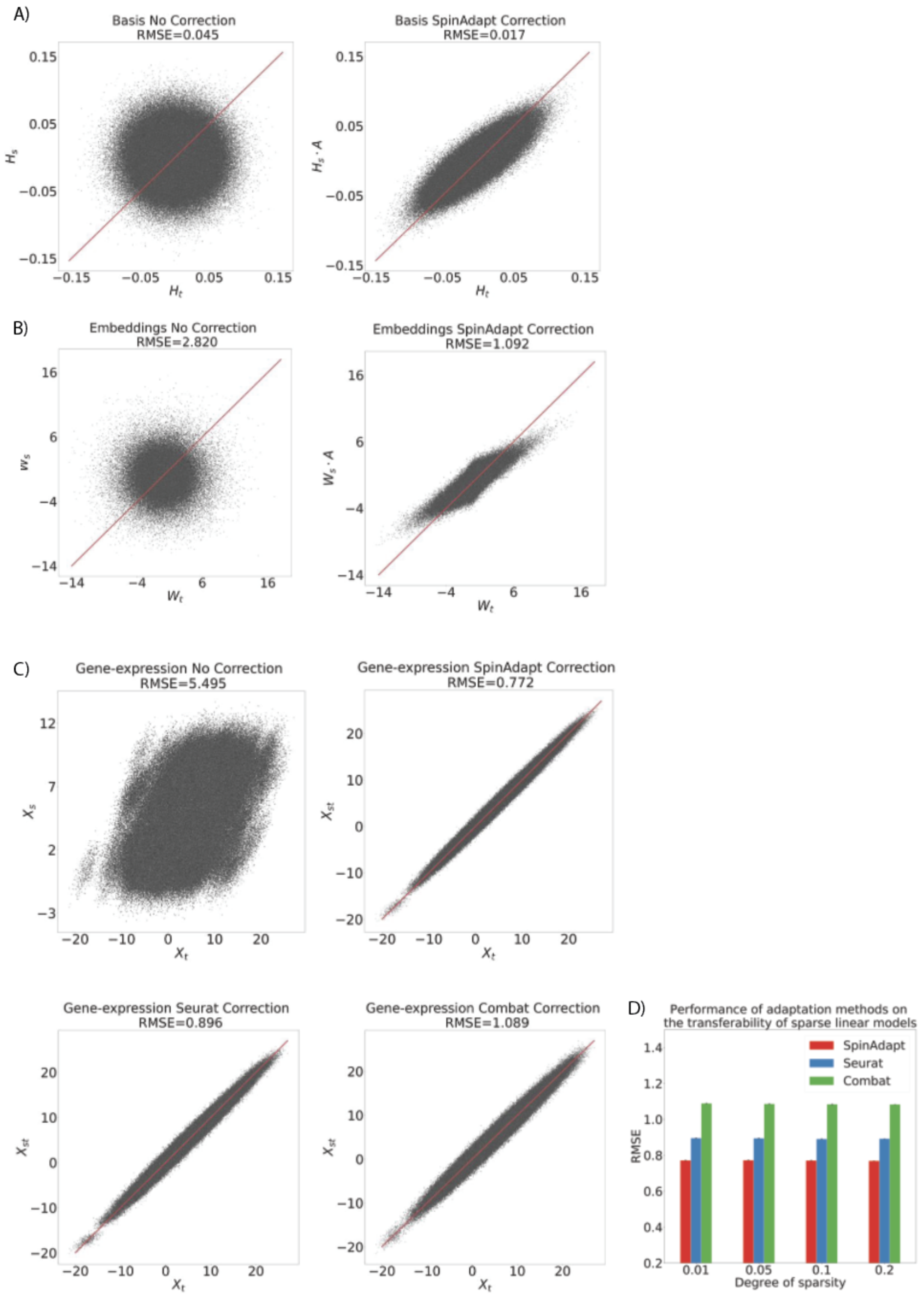


**Supplementary Figure 6.** Paired simulated data experiment. **A**) Scatter plots of the target basis and uncorrected source basis (RMSE = 0.045), and the target basis and the learned SpinAdapt corrected source basis (RMSE = 0.017). **B**) Scatter plots of target embedding and the uncorrected source embedding (RMSE = 2.82), and target embedding and SpinAdapt corrected source embedding (RMSE = 1.092). **C**) Scatter plots of paired expression values in the target and: un-corrected source (RMSE = 5.5) SpinAdapt corrected source (RMSE = 0.77), Seurat corrected source for Seurat (RMSE = 0.89), and Combat corrected source (RMSE = 1.09). **D**) Performance of 500 randomly generated sparse linear models evaluated on simulated target and on corrected source. Experiment repeated for each method across varying sparsity levels. SpinAdapt outperforms Seurat and ComBat for each sparsity level based on RMSE.

#### Supplementary Tables

| Cancer type | | Source | | Target | |
|---|---|---|---|---|---|
| Bladder cancer | Dataset | Seiler | | TCGA | |
| | Assay platform | Affy HumanExon 1.0 ST | | RNA-Seq | |
| | Subtype | Samples | % | Samples | % |
| | Ba-Sq | 114 | 38% | 116 | 40% |
| | LumP | 76 | 25% | 88 | 31% |
| | LUMU | 69 | 23% | 37 | 13% |
| | Stroma-rich | 24 | 8% | 31 | 11% |
| | LumNS | 18 | 6% | 16 | 6% |
| | Total | 301 | 100% | 288 | 100% |
| Colorectal cancer | Dataset | GSE14333 | | TCGA | |
| | Assay platform | Affymetrix hgu-133plus2 | | RNA-Seq | |
| | Subtype | Samples | % | Samples | % |
| | CMS1 | 21 | 16% | 62 | 19% |
| | CMS2 | 55 | 43% | 119 | 37% |
| | CMS3 | 21 | 16% | 50 | 16% |
| | CMS4 | 31 | 24% | 89 | 28% |
| | Total | 128 | 100% | 320 | 100% |
| Breast cancer | Dataset | TCGA | | SCAN-B | |
| | Assay platform | RNA-Seq | | RNA-Seq | |
| | Subtype | Samples | % | Samples | % |
| | Basal | 166 | 18% | 206 | 11% |
| | Her2 | 73 | 8% | 207 | 11% |
| | LumA | 470 | 52% | 1059 | 55% |
| | LumB | 192 | 21% | 466 | 24% |
| | Total | 901 | 100% | 1938 | 100% |
| Pancreatic cancer | Dataset | TCGA | | Bailey | |
| | Assay platform | RNA-Seq | | RNA-Seq | |
| | Subtype | Samples | % | Samples | % |
| | ADEX | 33 | 27% | 9 | 13% |
| | Immunogenic | 25 | 21% | 20 | 29% |
| | Progenitor | 42 | 35% | 21 | 30% |
| | Squamous | 21 | 17% | 20 | 29% |
| | Total | 121 | 100% | 70 | 100% |

**Supplementary Table 1:** Datasets used in the study.

| Cancer type | Subtype | SpinAdapt | Seurat | ComBat | Limma | Scanorama |
|---|---|---|---|---|---|---|
| Bladder cancer | Ba-Sq | 0.98 (8e-4) | 0.90 (5e-3) | 0.94 (1e-3) | X | X |
| | LumNS | 0.53 (9e-3) | 0.37 (2e-2) | 0.27 (9e-3) | X | X |
| | LumP | 0.74 (3e-3) | 0.65 (6e-3) | 0.62 (3e-3) | X | X |
| | LumU | 0.66 (4e-3) | 0.63 (1e-2) | 0.37 (4e-3) | X | X |
| | Stroma-rich | 0.79 (7e-3) | 0.44 (2e-2) | 0.24 (1e-2) | X | X |
| Colorectal cancer | CMS-1 | 0.78 (5e-3) | 0.70 (8e-3) | 0.44 (6e-3) | X | X |
| | CMS-2 | 0.93 (2e-3) | 0.89 (3e-3) | 0.83 (4e-3) | X | X |
| | CMS-3 | 0.85 (6e-3) | 0.45 (1e-2) | 0.26 (8e-3) | X | X |
| | CMS-4 | 0.86 (3e-3) | 0.87 (6e-3) | 0.84 (3e-3) | X | X |
| Breaset cancer | Her2 | 0.78 (3e-3) | 0.76 (4e-3) | 0.68 (2e-3) | X | X |
| | LumA | 0.91 (6e-4) | 0.86 (1e-3) | 0.91 (5e-4) | X | X |
| | LumB | 0.77 (2e-3) | 0.72 (2e-3) | 0.76 (1e-3) | X | X |
| | Basal | 0.99 (5e-4) | 0.98 (7e-4) | 0.97 (6e-4) | X | X |
| Pancreatic cancer | ADEX | 0.81 (3e-3) | 0.49 (1e-2) | 0.54 (6e-3) | X | X |
| | Progenitor | 0.86 (3e-3) | 0.65 (7e-3) | 0.69 (4e-3) | X | X |
| | Immunogenic | 0.64 (7e-3) | 0.40 (7e-3) | 0.49 (8e-3) | X | X |
| | Squamous | 0.72 (5e-3) | 0.72 (6e-3) | 0.76 (4e-3) | X | X |

**Supplementary Table 2.** Average F-1 scores for tumor subtype prediction: mean (std error).

| | | SpinAdapt | Seurat | ComBat | Limma | Scanorama |
|---|---|---|---|---|---|---|
| Bladder cancer | tLISI | 1.6 (3e-2) | 1.62 (3e-2) | 1.73 (3e-2) | 1.68 (3e-2) | 1.57 (3e-2) |
| | bLISI | 1.85 (7e-3) | 1.88 (5e-3) | 1.60 (1e-2) | 1.68 (1e-2) | 1.74 (1e-2) |
| | bLISI / tLISI | 1.35 (2e-2) | 1.33 (2e-2) | 1.13 (2e-2) | 1.20 (2e-2) | 1.30 (2e-2) |
| | Silhouette | 0.16 (1e-2) | 0.16 (1e-2) | 0.14 (1e-2) | 0.14 (1e-2) | 0.19 (1e-2) |
| Colorectal cancer | tLISI | 1.33 (2e-2) | 1.40 (2e-2) | 1.50 (3e-2) | 1.53 (3e-2) | 1.50 (3e-2) |
| | bLISI | 1.66 (1e-2) | 1.63 (1e-2) | 1.43 (2e-2) | 1.48 (1e-2) | 1.58 (1e-2) |
| | bLISI / tLISI | 1.35 (2e-2) | 1.29 (2e-2) | 1.06 (2e-2) | 1.08 (2e-2) | 1.18 (2e-2) |
| | Silhouette | 0.32 (1e-2) | 0.28 (1e-2) | 0.22 (1e-2) | 0.20 (1e-2) | 0.24 (1e-2) |
| Breast cancer | tLISI | 1.41 (9e-3) | 1.45 (1e-2) | 1.44 (9e-3) | 1.45 (9e-3) | 1.51 (1e-2) |
| | bLISI | 1.63 (5e-3) | 1.68 (5e-3) | 1.59 (5e-3) | 1.62 (5e-3) | 1.46 (8e-3) |
| | bLISI / tLISI | 1.25 (7e-3) | 1.29 (8e-3) | 1.21 (7e-3) | 1.23 (8e-3) | 1.05 (7e-3) |
| | Silhouette | 0.22 (6e-3) | 0.21 (6e-3) | 0.19 (6e-3) | 0.17 (6e-3) | 0.11 (7e-3) |
| Pancreatic cancer | tLISI | 1.74 (4e-2) | 1.85 (5e-2) | 1.96 (4e-2) | 1.85 (5e-2) | 1.83 (5e-2) |
| | bLISI | 1.82 (1e-2) | 1.79 (2e-2) | 1.76 (2e-2) | 1.75 (2e-2) | 1.72 (2e-2) |
| | bLISI / tLISI | 1.12 (2e-2) | 1.06 (2e-2) | 0.97 (2e-2) | 1.03 (2e-2) | 1.02 (2e-2) |
| | Silhouette | 0.22 (2e-2) | 0.20 (2e-2) | 0.14 (2e-2) | 0.18 (2e-2) | 0.19 (2e-2) |

**Supplementary Table 3.** Integration metrics reported across the 4 cancer datasets: mean (std error).

tLISI: Tissue LISI. Lower values indicate subtype clusters are more homogenous.

bLISI: Batch LISI. Higher values indicate more homogenous mixing of the two datasets.

bLISI / tLISI: Ratio of Batch LISI to tLISI. Higher values indicate more homogenous mixing of the two datasets or/and homogenous subtype clusters.

Silhouette: Silhouette Score. Higher values indicate subtype clusters are more homogenous.

| Cancer type | Subtype | SpinAdapt-Seurat | SpinAdapt-ComBat | Seurat-ComBat |
|---|---|---|---|---|
| Bladder cancer | Ba-Sq | 9.90E-05 | 1.60E-02 | 3.60E-02 |
| | LumNS | 2.50E-01 | 3.90E-02 | 4.50E-01 |
| | LumP | 4.40E-01 | 4.80E-07 | 8.80E-07 |
| | LumU | 4.20E-01 | 1.90E-06 | 1.10E-05 |
| | Stroma-rich | 2.00E-03 | 1.20E-04 | 2.20E-01 |
| Colorectal cancer | CMS-1 | 2.30E-01 | 3.90E-03 | 3.10E-02 |
| | CMS-2 | 4.50E-01 | 3.90E-03 | 9.40E-02 |
| | CMS-3 | 2.70E-03 | 1.20E-04 | 2.50E-01 |
| | CMS-4 | 1.70E-01 | 6.20E-02 | 6.20E-01 |
| Breaset cancer | Her2 | 5.50E-01 | 3.70E-04 | 1.70E-02 |
| | LumA | 1.30E-12 | 7.40E-01 | 3.50E-12 |
| | LumB | 6.70E-01 | 2.10E-01 | 2.20E-01 |
| | Basal | 5.00E-01 | 7.00E-02 | 2.20E-01 |
| Pancreatic cancer | ADEX | 6.10E-05 | 2.40E-04 | 8.80E-01 |
| | Progenitor | 3.40E-03 | 2.00E-03 | 6.20E-01 |
| | Immunogenic | 3.70E-03 | 1.60E-02 | 6.20E-01 |
| | Squamous | 5.00E-01 | 2.50E-01 | 1.00E+00 |

**Supplementary Table 4.** P values for comparing prediction performance between methods for tumor subtype prediction, reported as median over 30 experiments.

| | | SpinAdapt-Seurat | SpinAdapt-ComBat | SpinAdapt-Limma | SpinAdapt-Scanorama |
|---|---|---|---|---|---|
| Bladder cancer | tLISI | 4.50E-06 | 9.50E-10 | 2.30E-07 | 9.60E-01 |
| | bLISI | 1.60E-02 | 9.00E-61 | 4.50E-34 | 1.50E-14 |
| | Silhouette | 6.50E-01 | 9.20E-06 | 3.60E-08 | 1.20E-05 |
| Colorectal cancer | tLISI | 2.30E-07 | 5.70E-17 | 5.20E-19 | 2.90E-17 |
| | bLISI | 1.10E-01 | 6.60E-31 | 1.10E-25 | 1.50E-06 |
| | Silhouette | 8.30E-05 | 4.50E-26 | 1.20E-26 | 2.90E-19 |
| Breast cancer | tLISI | 9.10E-13 | 2.30E-15 | 8.70E-15 | 1.70E-42 |
| | bLISI | 1.20E-30 | 4.00E-12 | 4.30E-01 | 5.20E-94 |
| | Silhouette | 2.00E-11 | 1.10E-129 | 1.30E-122 | 7.00E-95 |
| Pancreatic cancer | tLISI | 2.20E-02 | 3.10E-10 | 1.60E-03 | 1.40E-02 |
| | bLISI | 3.20E-02 | 3.70E-05 | 1.60E-04 | 8.90E-08 |
| | Silhouette | 1.30E-01 | 8.10E-06 | 7.30E-03 | 4.50E-02 |

**Supplementary Table 5.** P values for comparing integration performance between SpinAdapt and other methods.

| Cancer type | | Source | Target |
|---|---|---|---|
| Colorectal cancer | Dataset | GSE14333 | TCGA |
| | Assay platform | Affymetrix hgu-133plus2 | RNA-Seq |
| | Subtype | 226 | 579 |
| Breast cancer | Dataset | TCGA | SCAN-B |
| | Assay platform | RNA-Seq | RNA-Seq |
| | Subtype | 1038 | 2919 |
| Pancreatic cancer | Dataset | PACA-AU | PACA-CA |
| | Assay platform | Array-based gene expression | RNA-Seq |
| | Samples | 267 | 186 |

**Supplementary Table 6:** Datasets used in the study.

| | | C-index | P-value | Hazard Ratio (HR) (95% CI for HR) |
|---|---|---|---|---|
| Colorectal cancer | SpinAdapt | 0.63 | 3.75e-02 | 4.24 (0.96-18.6) |
| | Seurat | 0.51 | 6.28e-01 | 0.85 (0.43-1.66) |
| | ComBat | 0.56 | 1.12e-01 | 0.57 (0.28-1.15) |
| Breast cancer | SpinAdapt | 0.66 | 9.72e-06 | 3.2 (1.86-5.50) |
| | Seurat | 0.59 | 1.06e-02 | 1.95 (1.16-3.29) |
| | ComBat | 0.61 | 2.48e-04 | 2.4 (1.48-3.89) |
| Pancreatic cancer | SpinAdapt | 0.65 | 1.74e-06 | 3.51 (2.04-6.04) |
| | Seurat | 0.62 | 1.99e-04 | 2.66 (1.56-4.54) |
| | ComBat | 0.62 | 6.48e-05 | 2.64 (1.61-4.32) |

**Supplementary Table 7.** Concordance index (C-index), P value, and Hazard Ratio (HR) with 95% Confidence Interval for comparing transfer of COX regression models using SpinAdapt, Seurat, and ComBat.

#### Supplementary note

**Privacy.** SpinAdapt is a batch correction method, which learns corrections between latent space representations that de-identify the sample-level information in each dataset. In this study, the PCA basis are chosen as the latent space representations for each dataset. SpinAdapt only requires access to data factors of each dataset for computation and application of dataset corrections, where the data factors consist of the PCA basis, gene-wise means, and gene-wise variances of each dataset. Therefore, for the transfer of an RNA model from a training dataset to a validation dataset, only the data factors of the training dataset need to be transferred along the RNA model, thereby maintaining ownership of the training dataset.

Next, we show that the data factors are privacy-preserving, since an expression dataset cannot be recovered from the PCA basis of the dataset. Let $X$ be an expression matrix of size {genes x samples}, let the columns of a matrix $W$ contain the eigenvectors of $XX^T$, let the columns of a matrix $V$ contain the eigenvectors of $X^TX$, and let $\Sigma$ be a diagonal matrix such that the entries are square roots of the eigenvalues of $XX^T$. Note that $W$ and $V$ are PCA basis for the columns and rows of the expression matrix $X$, respectively. An SVD decomposition of the expression matrix $X$ can be written as $X = W \Sigma V^T$, which implies $W = X V\Sigma^{-1}$, since the columns of matrix $V$ are orthonormal and $\Sigma$ is a diagonal matrix. Let $H$ be another matrix $H = V\Sigma^{-1}$. Then, $W = X V\Sigma^{-1}$can be rewritten as $W = X H$. From $W = X H$, it can be seen that the PCA basis matrix $W$ is obtained via an affine transformation $H$, which de-identifies the original expression matrix $X$. Such affine transformations are called matrix masks in the privacy literature [14]. Since the matrix mask $H$ is kept private, while only the PCA basis matrix $W$ is made public, recovering original sample-level data $X$ from $W$ involves solving a highly underdetermined system [15]. Hence, the affine transformation $H$ de-identifies the original data $X$, and the PCA factors $W$ are privacy-preserving representations of the expression data $X$.

**Simulation details.** Let us set p = 1000 and n = 500, where p is number of genes and n is number of patients. To evaluate SpinAdapt, we simulated n source samples (denoted as $X_s$) from a p-dimensional multivariate normal distribution with mean $\mu_s$ and covariance matrix $\Sigma_s := CC'$, where $\mu_s$ is a 1000-dimensional vector of uniform(0,10) random variables, C' is the transpose of the matrix C, and C is a p-by-p matrix of standard normal random variables. To model the dataset bias, we generate a p-by-p matrix B, which is equal to the sum of an identity matrix and a p-by-p matrix of standard normal random variables. We generated target data $X_t$ using $X_t = X_s B + \epsilon_t$, where $\epsilon_t$ is a n-by-n matrix of normal random variables. The target data $X_t$ represents n instances of a p-dimensional multivariate normal distribution with mean $\mu_s B$ and covariance matrix $\Sigma_t = B'\Sigma_s B + 0.25 I_{1000}$ .

We train SpinAdapt on the PCA basis of Xs and Xt, which are Hs and Ht, respectively:

$$W_s, H_s = PCA(X_s)$$

$$W_t, H_t = PCA(X_t).$$

The basis correction (A is a linear basis change operator for SpinAdapt in this setting) to Hs approximates Ht with low error (RMSE = 0.017, **Supplementary Figure 6A**). We correct the source embeddings through Ws*A, which estimates Wt with RMSE = 1.092 (**Supplementary Figure 6B**). We evaluated the performance of SpinAdapt on correction of the simulated biased source dataset $X_s$, drawing comparisons with no correction, Seurat, and Combat. To compare the correction performance for each method, we computed the RMSE for each technique. All of the methods achieved lower error than no correction (**Supplementary Figure 6C**). SpinAdapt outperformed both Seurat and Combat with an RMSE = 0.772 vs RMSE = 0.896 and RMSE = 1.089, respectively.

Testing the transferability of an arbitrary function across target and source corrected data, we generated 500 sparse linear models, each consisting of a normalized p-dimensional coefficient vector of standard normal random variables. We repeat the experiment for sparsity levels of 1%, 5%, 10%, and 20%, such that 10, 50, 100, and 200 components of the coefficient vectors have non-zero elements, respectively. For each sparsity level, we denote this collection of models as $\left\{\beta^{(m)}\right\}_{m=1}^{500}$. For each sparsity level, we calculated the RMSE of the differences between observed target values ($y_i^{(m)} = x_{t_i}\beta^{(m)}$) and predicted values for the corrected source data ($\hat{y}_i^{(m)} = x_{st_i}\beta^{(m)}$), where $x_{st_i}$ represents source corrected dataset (**Supplementary Figure 6D**). SpinAdapt outperforms Seurat and ComBat in terms of RMSE, for any chosen level of sparsity.
